# Large-scale culturing of the tree microbiome enables targeted disease suppression

**DOI:** 10.1101/2025.11.18.689053

**Authors:** Alejandra Ordonez, Marine C. Cambon, Jim Downie, Anparasy Kajamuhan, Usman Hussain, Bridget Crampton, Megan Richardson, Marcus Jones, Robert Green, Bethany Pettifor, Ed Pyne, Jasen Finch, Carrie Brady, Sandra Denman, James E. McDonald

**Author notes:** Corresponding authors: James McDonald and Alejandra Ordonez.

## Abstract

The tree microbiome is essential for host health and pathogen suppression. Synthetic microbial communities (SynComs) are emerging as important tools to understand microbiome dynamics and engineer microbiomes to harness beneficial properties. However, while the rational design, assembly and application of SynComs requires representative microbiota isolate collections combined with functional information, microbial culture collections from tree species such as oak (*Quercus*) are critically lacking. Here, we generated an oak microbiota culture collection comprising >30,000 isolates from 150 oak trees across Britain, belonging to key bacterial and fungal taxa that represented 61% of the total bacterial sequences and 87% of total fungal sequences as determined by culture-independent sequencing. Over 22,000 isolates were screened for suppression of bacterial species associated with degradation of live stem tissue in trees affected by Acute Oak Decline (AOD), identifying 341 bacterial isolates that suppressed oak pathogens. *In vitro* screening of 40 randomly assembled SynComs demonstrated that oak microbiota SynComs can suppress oak pathogenic bacteria associated with AOD. Inoculation of a disease-suppressive SynCom into the stem of oak seedlings and logs prior to pathogen challenge reduced the quantities of the bacteria, *Brenneria goodwinii* and *Gibbsiella quercinecans,* by 56% and 87%, respectively, in seedlings, and 71% and 95% in logs. This work demonstrates that the tree microbiome can be engineered using disease suppressive SynComs.

## Introduction

Trees possess taxonomically and functionally distinct microbiomes across different tissue types that are essential for host health through nutrient provision, immune priming, promoting stress tolerance and pathogen suppression (Mendes et al., 2011; Gu et al., 2020; Arnold et al., 2025). Tree microbiomes also influence global biogeochemical cycles (Jeffrey et al., 2021; Gauci et al., 2024), and their composition and function are influenced by environmental factors and the presence of disease (Baldrian, 2017; Cambon et al., 2025; Downie et al., 2025). As global tree species are increasingly impacted by changing climatic conditions and disease outbreaks, understanding the ecological factors underlying assembly and dynamics of the tree microbiome becomes an emerging research need (Baldrian et al., 2023; Downie et al., 2025).

Despite the importance of the tree microbiome in sustaining forest ecosystems and the ecological and economic services they provide, our understanding of tree microbiome dynamics is comparatively underexplored relative to marine, soil, human and plant microbiomes. While recent large-scale DNA sequencing studies have illuminated the complexity and ecological importance of tree microbiomes (Arnold et al., 2025; Downie et al., 2025), large-scale microbial culture collections are transforming our understanding of mechanistic interactions in other globally important microbiomes, including marine (Sanz-Sáez et al., 2020), human (Hitch et al., 2025; Huang et al., 2023) and model plant microbiomes (Bai et al., 2015; Figueroa-López et al., 2016; Durán et al., 2018), but have not been generated for tree species. The generation of large-scale microbial culture collections is therefore essential to enable physiological analysis of individual members of the tree microbiota and drive mechanistic understandings of tree microbiomes, for example using synthetic microbial communities (Cambon et al., 2025).

Synthetic microbial communities (SynComs) - microbial consortia designed to mimic natural microbiomes, or to perform a specific function - are emerging as promising systems to dissect microbiome dynamics and engineer microbiomes towards beneficial properties (Lawson et al., 2019; Emmenegger et al., 2023; Pfeilmeier et al., 2024), and have been utilised in both host-associated and environmental contexts for bioremediation, chemical engineering, growth promotion, and disease suppression in animal, plant and human diseases (Mehlferber et al., 2024). SynComs have transformed our understanding of microbiome assembly and dynamics in plants (Vorholt et al., 2017; Vogel et al., 2021) and have been harnessed to improve crop performance (Kaur et al., 2022) and suppress pathogens (Zhou et al., 2022). However, most advances in plant protective engineered microbiomes have been conducted in model plants like *Arabidopsis thaliana* (Emmenegger et al., 2023; Pfeilmeier et al., 2024; Ryback et al., 2022) and crop plants (Zhou et al., 2022; Su et al., 2024; Virgo et al., 2025). SynComs also have significant potential to advance tree microbiome research; however, as trees can live for centuries, are large, and comprised of woody tissues, characterising and engineering tree microbiomes raises specific challenges (Addison et al., 2024; Cambon et al., 2025). In the phyllosphere, inoculation of mixed microbial consortia suppressed an oomycete leaf pathogen of cacao trees (Arnold et al., 2003), a fungal pathogen of poplar trees (Busby et al., 2016) and the causal agent of white pine blister rust (Ganley et al., 2008). Allsup et al. (2023) demonstrated that soil microbiome transplants form trees adapted to cold, warm and drought-stressed environments enhanced tolerance to these stressors. Together, these studies highlight the potential to modulate the tree microbiome.

Here, we generated an oak-associated microbiota collection comprising >30,000 isolates that were broadly representative of healthy oak trees across Britain and performed agar-based high-throughput functional screening of >22,000 isolates to identify bacteria that suppress *B. goodwinii* (Bg) and *G. quercinecans* (Gq), two bacterial species associated with degradation of live stem tissue in mature oak trees impacted by Acute Oak decline (AOD) (Denman et al., 2017). AOD is a complex decline disease affecting mature oak trees in Europe and beyond (Gosling et al., 2024), driven by environmental stressors (Brown et al., 2018), a shift in stem microbiome composition to a canker-causing microbial community (Denman et al., 2017), and larval activity of the bark-boring beetle *Agrilus biguttatus* that enhances pathogen virulence (Doonan et al., 2020; Cambon et al., 2023). Oak trees impacted by AOD can die within 3-5 years due to impaired vascular transport and crown dieback (Denman et al., 2014). Using our functionally screened isolate collection, we assembled random SynComs of varying species richness to validate the disease suppressive properties of assembled SynComs against Bg and Gq *in vitro*. Subsequently, we inoculated disease-suppressive SynComs into oak seedlings and logs and tested their efficacy for disease suppression in plants both before and after pathogen challenge. One SynCom applied to oak seedlings and logs prior to pathogen challenge had a protective effect against pathogen establishment, demonstrating the potential to engineer the tree microbiome to promote tree health and manage disease.

## Results

### Characterising an extensive oak microbiota culture collection

We isolated microorganisms from 450 leaf, stem and root/rhizosphere (here onwards referred as rhizosphere) samples collected from 150 oak trees across Britain, from asymptomatic trees on sites where AOD was present (AOD sites), nearby sites with no reports of AOD (Near-AOD sites) and sites outside the AOD predisposition zone (Far-AOD sites). We obtained 14,122 bacterial and fungal isolates using agar plating, of which, 4,973 were from rhizosphere tissue, 5,780 from leaf tissue and 3,368 from stem tissue. Isolation by dilution-to-extinction was performed by inoculating 86,400 wells in 96-well plates with diluted cell suspensions from leaf and rhizosphere tissue only, as we previously demonstrated that dilution-to-extinction was not necessary for isolation of stem microbiota (Ordonez et al., 2025). We produced 96-well plates where 30 – 50 % of the wells contained visible growth, representing either a pure isolate or a low-richness community (hereafter, D2E-derived cultures), corresponding to ∼12,960 bacterial and fungal cultures from leaf tissue and ∼12,960 cultures from rhizosphere samples. In total, this resulted in a microbial culture collection containing more than 30,000 isolates, 14,122 obtained by agar plating and ∼25,920 from D2E, from 150 asymptomatic oak trees.

We determined the taxonomic richness of all isolates obtained by agar plating (n = 150 96-well plates) and 360 of the dilution-to-extinction plates (out of 900; at least one replicate per site, tree and tissue type) by pooling all wells of each 96-well plate. Amplicon sequencing of the V4 region of the 16S rRNA gene and the ITS1 was performed for bacterial and fungal identification, respectively. From bacterial isolates, we obtained 1,233 ASVs, representing 27 orders and 134 genera, with *Burkholderiales* (260 ASVs, 21%), *Xanthomonadales* (180 ASVs, 15%), *Pseudomonadales* (167 ASVs, 14%) and *Enterobacterales* (131 ASVs, 10%), from phylum *Pseudomonadota*, and *Paenibacillales* from phylum *Firmicutes* (89 ASVs, 7%), emerging as the most abundant bacterial orders in the collection (Figure 1A). Sequencing of the ITS1 revealed 650 fungal ASVs, representing 57 orders and 165 genera. The most abundant taxa were *Tremellales* (phylum *Basidiomycetes*, 117 ASVs, 18%), *Hypocreales* (67 ASVs, 10%), *Eurotiales* (51 ASVs, 8%), *Capnodiales* (44 ASVs, 7%) and *Dothideales* (39 ASVs, 6%), all belonging to phylum *Ascomycetes* (Figure 1B). The composition of the microbial culture collection was significantly different across sites, AOD predisposition zones and isolation methods (Supplementary Table 1).

**Figure 1.**
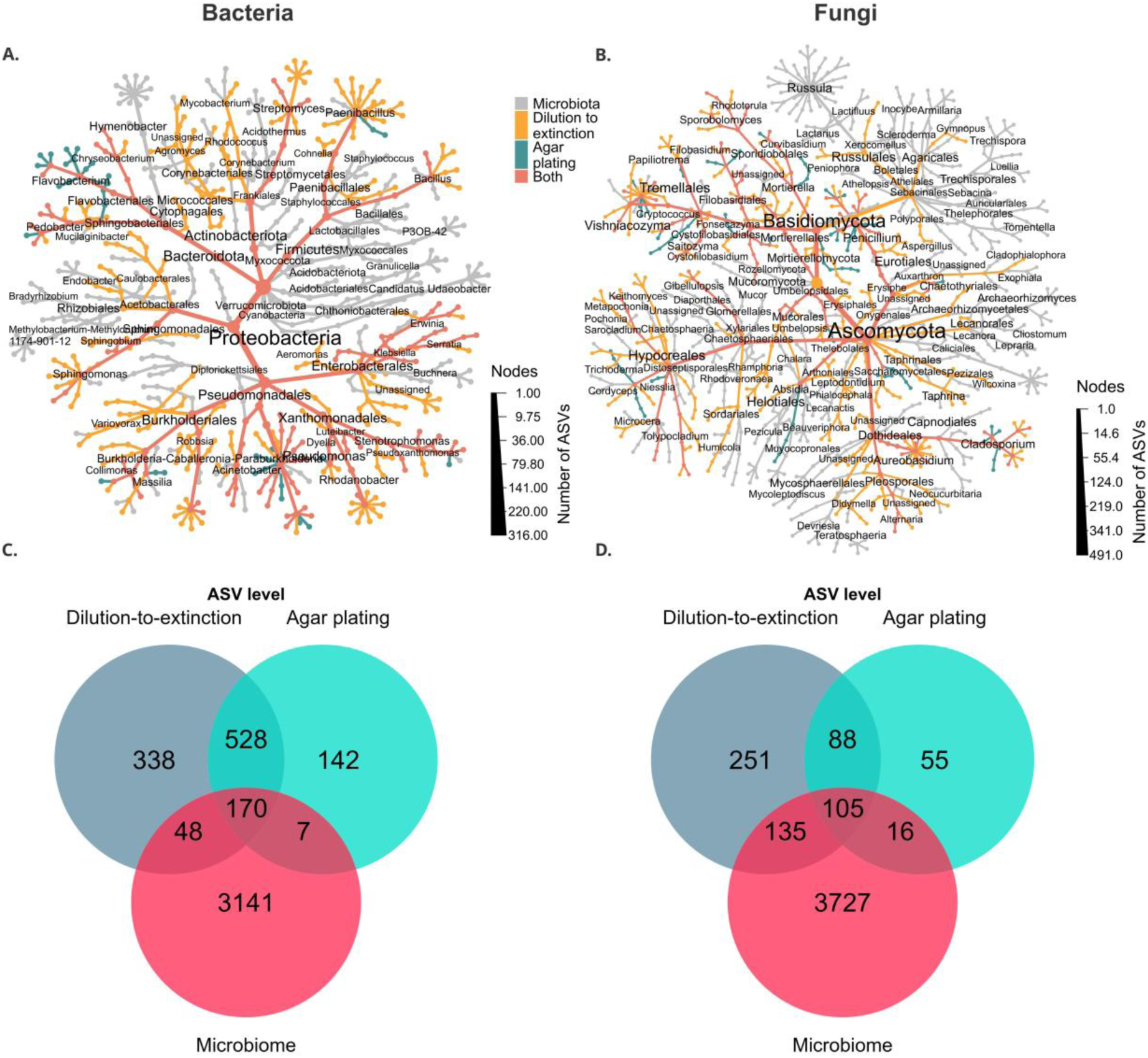
Comparison of the taxonomic richness and composition of a microbial culture collection obtained from the microbiota of 150 oak trees sampled across Great Britain and the oak microbial community analysed by culture-independent analysis. Oak microbial isolates obtained by agar plating and dilution-to-extinction were analysed by 16S rRNA gene and ITS community profiling for characterization of bacteria and fungi, respectively, and compared against the community profiles of the microbiota present in the same oak tissue samples (relative abundance > 0.1 % in at least one sample). A) and B) Number of bacterial and fungal ASVs, respectively, detected in oak microbial isolates and D2E-based cultures (pure isolates and low-richness communities obtained by dilution-to-extinction) vs number of ASVs detected by culture-independent analysis. C) and D) Taxonomic composition of bacterial and fungal communities and isolates/D2E-cultures of the oak microbiota at phylum, class, order and genus levels. Taxa obtained from culture-independent analysis are depicted in grey colour (Microbiota), taxa captured by agar plating and dilution-to-extinction are depicted in green and yellow colour, respectively, and ASVs captured by both isolation methods are depicted in pink colour (both).

The culture collection contained bacterial isolates/D2E-cultures representing 33% of classes, 45% of orders, 49% of families, 39 % of genera and 16% of ASVs (RA > 0.1 %) identified via culture-independent sequencing of the oak microbiome of the same plant tissue samples (Figure 1C) (Downie et al., 2025). These included representatives of orders *Rhizobiales, Burkholderiales, Pseudomonadales, Sphingomonadales, Enterobacterales* and others, which collectively accounted for 61% of the sequencing reads in the culture-independent analysis (Figure 1A). Similarly, fungal isolates represented 76% of classes, 52% of orders, 39% of families, 25 % of genera and 11% of ASVs (relative abundance (RA) > 0.1%) detected by culture-independent sequencing in the same samples (Figure 1D). Notable orders included *Dothideales, Russulales, Filobasidiales, Capnodiales, Erysiphales* and others, representing 87% of the reads obtained by culture-independent analysis (Figure 1B).

Taxa represented in Figures 1A and 1B include some groups obtained from microbial isolations that were absent in the culture-independent analysis. 1,008 bacterial and 394 fungal ASVs, were exclusively detected from isolates/D2E-cultures, and the most abundant ones (RA > 0.1% in at least one sample) were assigned to 8 additional bacterial families and 31 fungal families. In contrast, 24 bacterial and 52 fungal families found in the culture-independent analysis of the oak microbiome were not represented in the culture collection (Extended Data 1). This suggests that while microbial isolations can enrich and allow detection of some taxa absent from culture-independent analysis, targeted isolation methods are required to capture misrepresented taxa and ensure higher coverage of the natural microbiome (Ordonez et al., 2025).

Our results overall demonstrate that a combination of different microbial isolation methods applied to the landscape scale captured a representative proportion of the taxonomic richness in the natural oak microbiome, as well as its variations at a geographical scale. Microbial culture-collections derived from model plant species such as *Arabidopsis thaliana* (Bai et al., 2015) have provided a foundational framework for subsequent plant microbiome research, enabling detailed investigations into microbiome assembly, dynamics and interactions (Schäfer et al., 2022; Vogel et al., 2021). The oak microbial culture collection generated in this study represents the largest and most comprehensive plant-associated culture collection generated to date, comprising bacterial and fungal taxa that are observed across a broad range of other plant and tree species (Pettifor & McDonald, 2021). Consequently, the culture collection represents a valuable resource to the wider scientific community, driving future research discoveries on plant microbiome dynamics and interactions (see Supplementary Table 2, which presents a database of the culture collection).

### Identification of microbiota associated with healthy oak trees

Because microbiota associated with tree health confer beneficial properties other than pathogen suppression, such as immune priming and growth promotion (Ryback et al., 2022; Emmenegger et al., 2023), we identified a subset of bacterial isolates in the culture collection that were associated with the healthiest trees in our cohort of 150 trees. Using the phenotypic decline index (PDI) of each tree as a proxy for tree health status, five trees with the lowest PDI and represented by the highest taxonomic richness in the culture collection were selected as sources of oak microbiota (hereafter, health-associated isolates). Using almost full-length 16S rRNA gene sequencing, we identified 66 species belonging to 33 genera, 23 families and 10 orders (n = 320 isolates, Figure 2A). In contrast, 46 species were represented by one or two isolates only (Extended Data 2). Of these isolates, 42 of the 66 species exhibited plant growth-promoting traits in previous studies, whereas 10 species were associated with plant and soil microbiomes, but no particular plant growth-promoting capability was documented (Supplementary Table 3).

**Figure 2.**
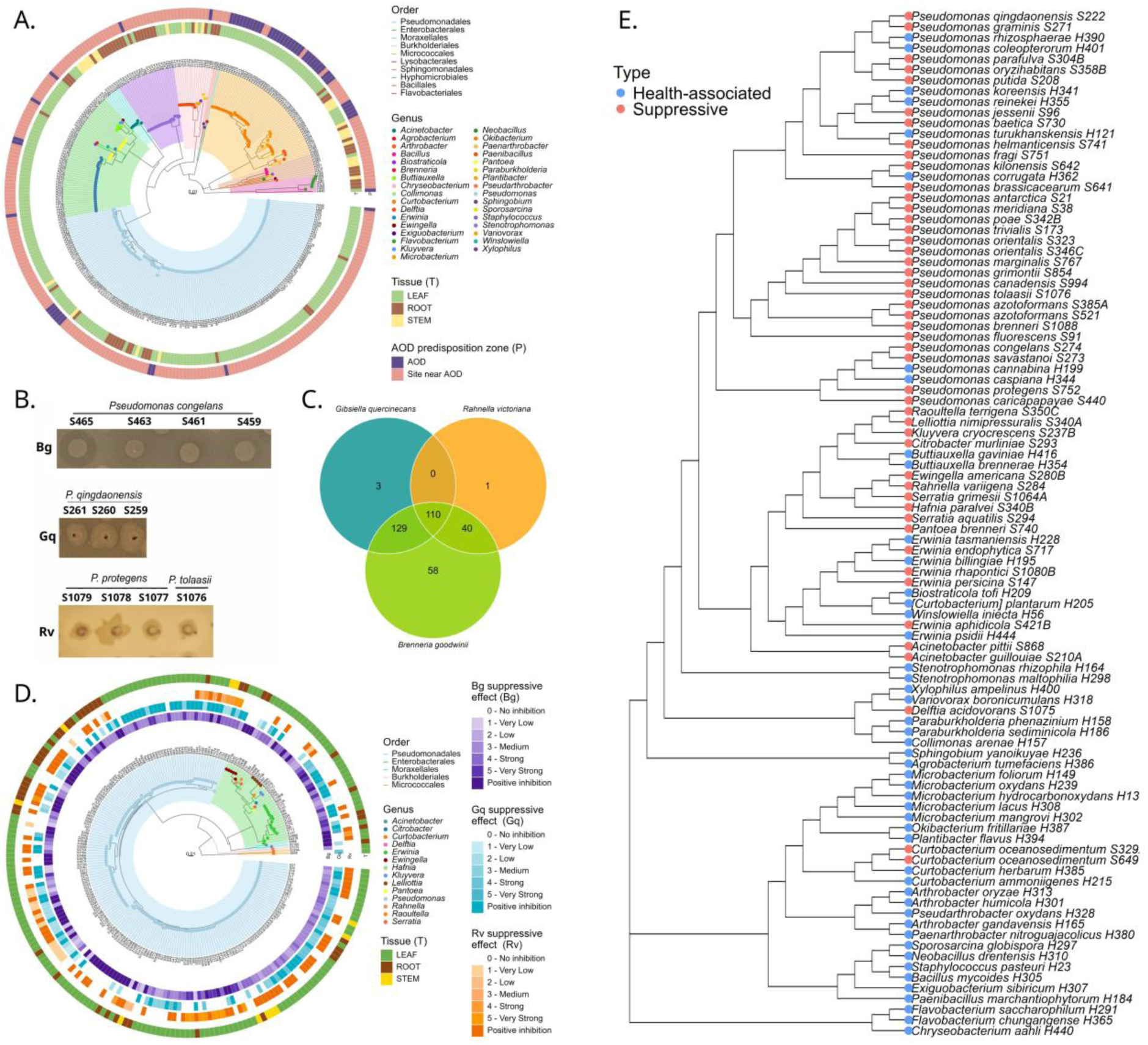
Identification of health-associated and suppressive isolates. A) Phylogenetic tree of microbial isolates obtained from oak trees exhibiting the lowest Phenotypic Decline Index amongst all the trees sampled (n = 150). B) Inhibition halos of different species of Pseudomonas against Brenneria goodwinii (Bg), Gibbsiella quercinecans (Gq) and Rahnella victoriana (Rv). C) Number of oak microbial isolates that exhibited in vitro suppressive activity against at least one of the members of the AOD pathobiome (Brenneria goodwinii, Gibbsiella quercinecans and Rahnella victoriana). D) Phylogenetic tree of the members of the oak microbiota with suppressive activity against the AOD pathobiome. The three first outer rings represent the suppressive effect of each isolate towards each bacterium in the AOD pathobiome (Bg: B. goodwinii, Gq: G. quercinecans and Rv: R. victoriana. E) Cladogram of 96 bacterial strains identified as health-associated or pathogen suppressive and used to prepare SynComs. Taxonomic assignment and phylogenetic trees were based on almost full-length sequences of the 16S rRNA gene obtained by Sanger sequencing technology.

### Screening for pathogen-suppressive oak microbiota

To identify members of the oak microbiota with suppressive activity against three bacterial species associated with degradation of live stem tissue in AOD, we screened approximately 22,762 bacterial and fungal isolates of the oak microbial culture collection (14,122 agar isolates, and one third of D2E-derived cultures, n = 300 x 96-well plates) against Bg, Gq, and the environmentally widespread bacterium, *Rahnella victoriana*, via detection of pathogen growth inhibition halos in dual agar-based culture assays. We identified 341 bacterial isolates with suppressive activity against at least one of the AOD-associated bacteria (hereafter, suppressive isolates) (Figure 2C); 138 isolates were obtained from AOD-Near sites, 117 from AOD-Far sites and 86 from AOD sites (Figure 2D). We obtained the full-length 16S rRNA gene of 225 of the isolates, revealing 47 species belonging to 14 genera, eight families and five orders (Figure 2D). The genus *Pseudomonas* was the most represented, with 28 species and 183 isolates, followed by *Erwinia* with four species and 16 isolates. Contrastingly, 24 other species were represented by only one isolate, and six species were represented by only two or three isolates (Extended Data 2). Of these, 11 species have been previously documented as pathogen suppressors or biocontrol agents (Supplementary Table 3), whilst 18 species had limited reports on pathogen-suppressiveness, but had been associated with other plant growth-promoting traits production of antimicrobials, hormones, siderophores and nutrient mobilization (Supplementary Table 3).

Interestingly, different isolates of the same species exhibited contrasting suppressive profiles, by either presenting inhibition halos of distinctive characteristics (i.e. size, transparency), or inhibiting different AOD-associated bacteria (Figure 2B). This was the case for 18 species, and mostly notable in isolates of *Pseudomonas marginalis* and *P. helmanticensis*, which consistently suppressed Bg, while some also inhibited Gq and/or *R. victoriana* (Figure 2D). This suggests functional differences at the strain level within the culture collection, which highlights the relevance of conducting large-scale isolation studies.

Overall, the collection of suppressive and health-associated microbial isolates comprised 96 species (49 health-associated, 30 suppressive and 17 that were both health-associated and suppressive, Extended Data 2), classified in 39 genera, 10 orders, six classes and four phyla (Figures 2A and 2D). For subsequent SynCom experiments, 96 bacterial strains representing 93 bacterial species were selected to assemble SynComs (Figure 2E).

### Randomly assembled SynComs suppress *B. goodwinii* and *G. quercinecans in vitro*

From our collection of 96 suppressive or health-associated isolates, we randomly assembled 40 SynComs with distinctive richness (8-96 members) and taxonomic composition (8-member SynComs, n = 11; 16-member SynCom, n = 7; 24-member SynCom, n = 7; 48-member SynCom, n = 7; 72-member SynCom, n = 7; 96-member SynCom, n = 1) and tested their suppressive effect against fluorescently labelled *B. goodwinii* FRB 186 + pBbE1K + Kan (RFP-Bg) and *G. quercinecans* BH1/65b + pBBRI-GFP + Kan (GFP-Gq), in microplate-based liquid co-culture assays. RFP-Bg and GFP-Gq were tested separately in SynCom inhibition assays where an inoculum composed of either bacterium combined with one SynCom in a 1:1 ratio was inoculated (10 % v/v) in a single well of a microplate. The OD_600_ of RFP-Bg and GFP-Gq was normalized before the assays, and all SynCom members were grown for 48 h and mixed in equal ratios, this ensured that Bg, Gq and each SynCom had approximately the same abundance in each assay.

Most SynComs prevented or reduced the growth of RFP-Bg and GFP-Gq compared with monoculture controls (Figure 3A and 3B). Overall, SynComs were more effective at suppressing GFP-Gq than RFP-Bg. Suppression indexes were calculated relative to the fluorescence of monoculture controls (See Supplementary Methods). Negative suppression indexes closer to −100 indicated stronger suppression (i.e. cell death or inability to express the fluorescent protein), and positive values closer to 100 indicated no suppression. For RFP-Bg, only half of the SynComs inhibited the growth of RFP-Bg (suppression indexes −86 to −1, Figure 3C, Extended Data 3), whilst the other half allowed the growth of RFP-Bg, although in a lesser extent compared to the RFP-Bg monoculture control (suppression indexes 4 – 63). In contrast, all SynComs were able to inhibit the growth of GFP-Gq (suppression indexes −69 to −14; Figure 3C, Extended Data 3).

**Figure 3.**
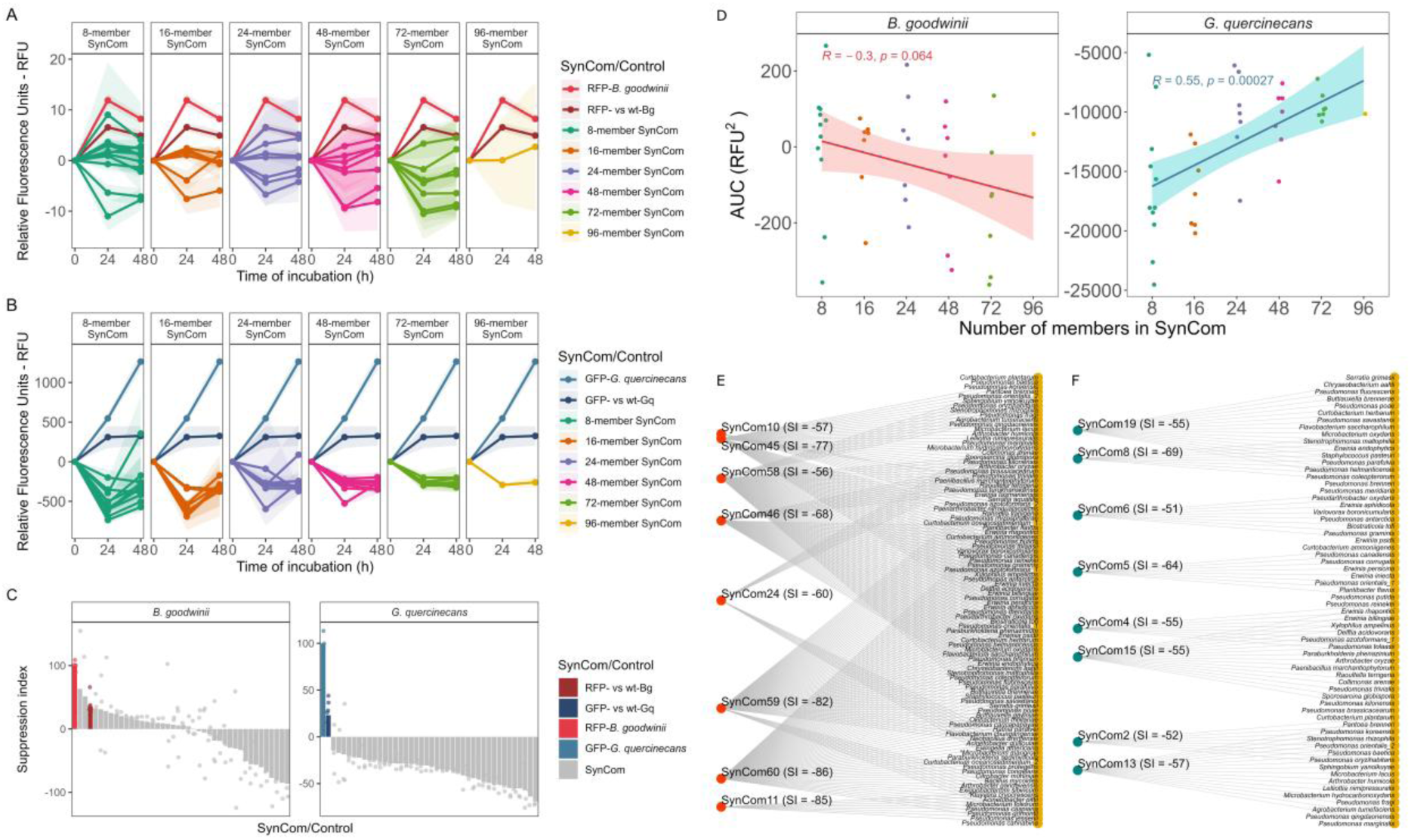
In vitro suppression of RFP-labelled B. goodwinii and GFP-labelled G. quercinecans by randomly assembled SynComs. A and B) Growth curves of RFP-Bg and GFP-Gq, respectively, in co-cultures with SynComs composed of different number of strains, C – D) Suppression index of SynComs compared with monocultures of RFP-Bg and GFP-Gq, respectively, and co-cultures of these strains with the type strains of Bg and Gq, respectively. Bars represent the mean suppression index, while dots indicate variation between replicates (n = 3 per SynCom). E – F) Linear regression between the number of strains in each SynCom and the area under the curve (AUC) calculated from fluorescence emission by RFP-Bg and GFP-Gq, respectively. Spearman correlation coefficient and p-values are displayed for each bacterium. G – H) Bipartite-style network plots representing the top 8 SynComs with highest pathogen suppressive activity against RFP-Bg and GFP-Gq, respectively, along with the identity of the strains in each SynCom. The suppression index (SI) is displayed in brackets next to each SynCom ID.

Interestingly, SynCom richness had contrasting effects on the growth of RFP-Bg and GFP-Gq; RFP-Bg growth was negatively correlated with SynCom richness based on area under curve (AUC) analysis (Spearman correlation R = −0.3, p-value = 0.06, Figure 3D) whereas GFP-Gq growth had a positive correlation with SynCom richness (Spearman R = 0.55, p-value < 0.001, Figure 3D). Higher diversity tends to be more resistant to invaders given than they tend to saturate available niches and utilize resources more efficiently than invaders (Hodgson et al., 2002; Spragge et al., 2023; Dooley et al., 2024). However, opposite trends have been previously reported in *Pseudomonas* populations (Becker et al., 2012; Mehrabi et al., 2016). This suggests that inherent or unique abilities of each bacterium can be shaping their interactions with other strains, rather than intracommunity dynamics themselves, for instance, differential sensitivity to antimicrobials, distinct resource preferences and varying competitive and fitness abilities of Bg and Gq (Hibbing et al., 2010; Brady et al., 2022).

A Generalized Linear Model (GLM) indicated a negative relationship between SynCom richness and RFP-Bg growth (p-value = 0.06) and a positive relationship with GFP-Gq growth (p-value < 0.01, Supplementary Table 4). To identify individual strains in SynComs associated with pathogen inhibition, we fitted GLMs including both SynCom richness and the presence/absence of each of the 96 bacterial species (Supplementary Table 4). The models identified a significant negative effect of *Stenotrophomonas maltophilia* H298 on the growth of GFP-Gq, a plant growth promoting bacteria in other plant systems (Rojas-Solís et al., 2018; Alexander et al., 2019), while none of the strains had a significant effect on growth of RFP-Bg.

### SynCom inoculation suppresses *B. goodwinii* and *G. quercinecans* in oak seedlings and logs

As many of the 40 SynComs tested *in vitro* had strong suppressive effects on RFP-Bg and/or GFP-Gq, and SynCom richness had contrasting impacts on pathogen suppression, ordination of the suppressive properties of the tested SynComs failed to identify a single SynCom that had superior suppressive properties. Consequently, for *in planta* SynCom inoculation aimed at targeted disease suppression, we decided to test three SynComs: SynCom1, containing 48 pathogen-suppressive strains (SynCom1); SynCom2, containing 48 health-associated strains, and SynCom3, containing all 96 strains (i.e. SynComs 1&2 combined). SynComs were inoculated into 2-year-old oak seedlings and logs from oak trees either ten days before or ten days after inoculation of the type strains of Bg and Gq (Bg:Gq inoculum ratio of 10:1). Swabs and microbial cultures were obtained at the end of the experiments (n = 10 inoculation points per treatment, Figure 4A). The pathogen load of Bg and Gq in the seedlings and logs trials was determined using species-specific *gyrB* and *rpoB* gene qPCR assays. For the seedling trial,16S rRNA gene community profiling was used to determine the relative abundance of AOD-derived OTUs (all OTUs assigned to Bg and Gq). For logs, we used 16S rRNA gene community profiling of both swabs and stem microbial cultures, to estimate the prevalence of Bg and Gq (percentage of samples where the bacteria were present, Extended Data 4). We observed significant positive correlations between the number of gene copies/mL detected by qPCR and the relative abundance of Bg and Gq from 16S rRNA gene profiling, highlighting the veracity of this approach to validate *in planta* SynCom inoculations (Extended Data 5).

**Figure 4.**
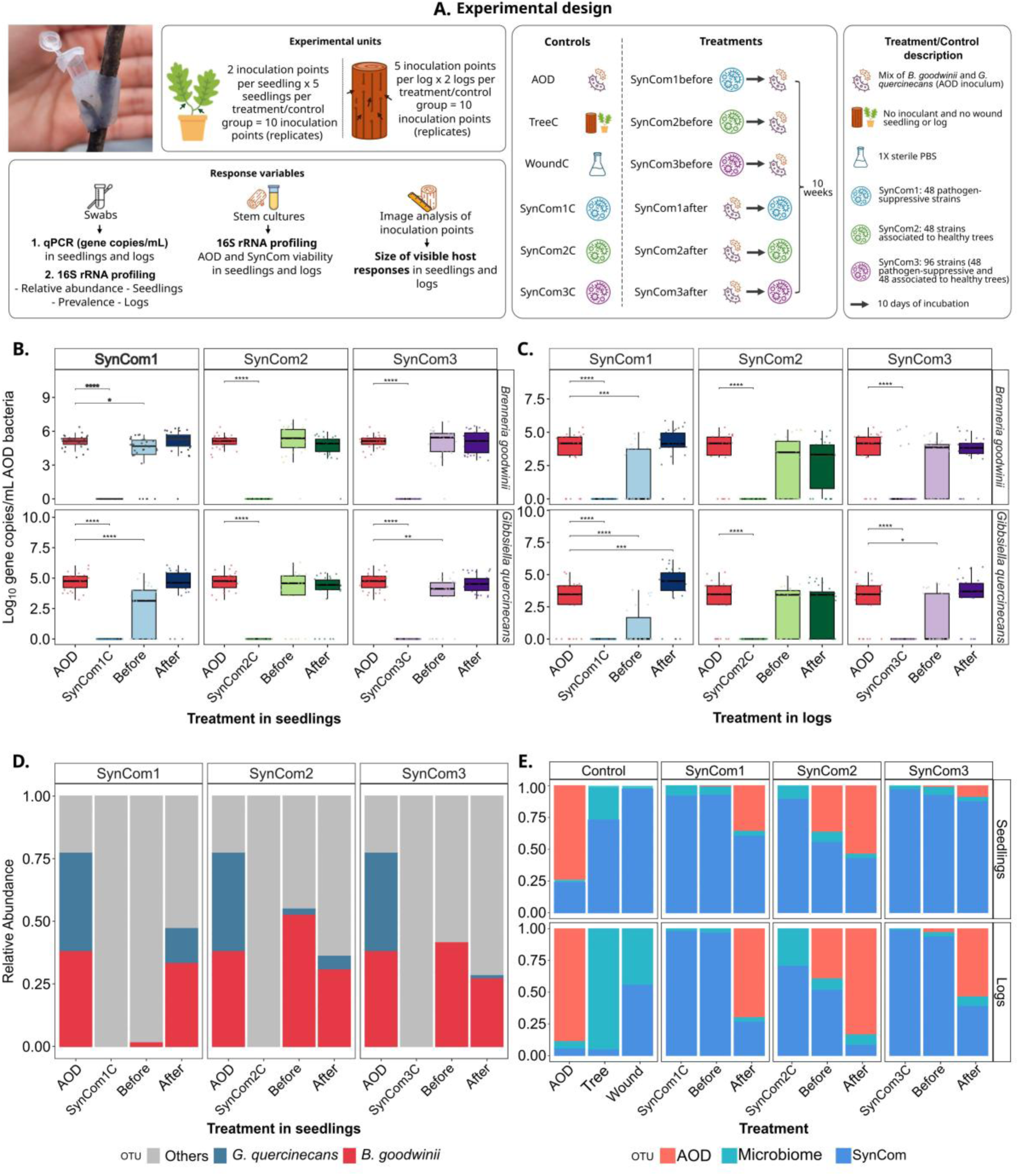
Effect of randomly assembled SynComs on the AOD-associated tree pathogens Brenneria goodwinii (Bg) and Gibbsiella quercinecans (Gq) in inoculation trials using oak seedlings and logs. A) Experimental design for testing the pathogen-suppressive activity of randomly assembled SynComs in oak seedlings and logs. B – C) Log_10_-transformed gene copies/mL of Bg and Gq detected in seedlings and logs, respectively. Wilcox rank-sum tests with Benjamini-Hochberg p-value adjustment were used to compare SynCom treatments against the AOD control. D) Relative abundance of Bg and Gq in seedlings trial, determined by microbiome profiling, E) Composition of microbial communities in stem cultures from oak seedlings and logs inoculated with Bg *and Gq and randomly assembled SynComs. The average relative abundance of each OTU category is represented per each treatment and control group*.

AOD inoculation control treatments exhibited high pathogen loads (Bg and Gq) in seedlings (1.3 – 2.5 x 10^5^ gene copies/mL and 64.4 ± 31.3% 16S rRNA gene relative abundance) and logs (1.5 – 3.0 x 10^4^ gene copies/mL, Figures 4B and 4C, respectively). Similarly, Bg and Gq were significantly enriched in AOD inoculation control treatments compared with controls that were not inoculated with the pathogens non-inoculated and non-wounded control (log2 fold change = 15.7 and 16.0, respectively, p-values < 0.05, Extended Data 6). We also checked visual symptoms and found that 7/10 replicates of the AOD control had symptoms in the seedlings trial (Extended Data 7), and that the size of the affected plant tissue was significantly higher in AOD control compared with non-inoculated control (Supplementary Table 5).

When SynComs were applied 10 days before inoculation of the pathogens, SynCom1 treatments had reduced gene copies/mL of Bg and Gq by 56 % and 87 % respectively in seedlings, and by 71 % and 95 % in logs, compared with the AOD controls (Figures 4B and 4C, respectively). This trend was also confirmed by 16S rRNA gene sequencing, where SynCom1 reduced the relative abundance of AOD-derived OTUs (Bg+Gq) by 94 % in seedlings (Figure 4D) and their prevalence by 82 % in logs (Extended Data 4). Similarly, SynCom3 (suppressive and health-associated isolates), when added in logs before pathogen inoculation reduced Bg and Gq gene copies/mL by 54% and 77%, respectively (Figure 4C), as well as their prevalence in 71 % (Extended Data 4). Consistently, SynCom3before also reduced Gq (76 % reduction of copy genes/mL and 26 % reduction of relative abundance of AOD-derived OTUs) in seedlings (Figures 4B and 4D). A differential abundance analysis in seedlings supported these results, indicating that Gq was significantly depleted when SynCom1 & 3 were added before pathogen inoculation (log2 fold change = −32.5 and −31.1, respectively, p-values < 0.05) (Extended Data 6).

Non-parametric Wilcox test showed significant differences in Bg and Gq gene copies/mL between the AOD control and treatments SynCom1before and SynCom3before in seedlings and logs (p-values ≤ 0.05, Figures 4B and 4C). Linear Mixed Effects Models (LMM), which account for potential pseudo-replication effects, confirmed significant reductions of Gq by SynCom1before in seedlings, and of both Bg and Gq by SynCom1before and SynCom3before in logs (p-values ≤ 0.05) (Extended Data 8). Generalized Linear Mixed Effect Models (GLMMs) were also applied, since LMM assumptions were not met (Extended Data 9), however, these did not show any significant differences in gene copies/mL (Extended Data 8). Nonetheless, GLMMs for relative abundance data (16S rRNA profiling) confirmed significant reductions of both bacteria in seedlings and logs by SynCom1before and in some cases by SynCom3before too (Extended Data 8). Altogether our results from *in planta* inoculation trials suggest that SynCom1 and SynCom3 applied before Bg and Gq significantly reduced the load of both bacteria, although, more replication is needed to increase the statistical power of analysis.

Inoculation of SynCom2 (health-associated isolates) before and after addition of Bg and Gq, as well as the application of SynCom1 and SynCom3 after did not result in a clear trend, with both increases and reductions occurring in Bg and Gq gene copies and 16S rRNA relative abundances compared to the AOD control. Interestingly, SynCom1 applied after Bg and Gq inoculation resulted in increased load of both AOD-associated bacteria in seedlings (73% and 21% increase of gene copies/mL of Bg and Gq, respectively) and in logs (over 2-fold increase of gene copies/mL of Bg and Gq, respectively). These results suggest that SynCom composition and the timing of SynCom application can have either positive or detrimental effects on pathogen suppression and highlights the importance of microbiota interactions in SynComs studies.

To confirm the viability of Bg, Gq and the SynComs inoculated into oak seedlings and logs, we cultured stem pieces from the inoculation points at the end of the experiment and used 16S rRNA gene sequencing of cultures to identify Bg, Gq and SynCom members, in addition to the endogenous oak seedling/log microbiota. 16S rRNA gene profiling revealed that Bg and Gq were culturable in all treatments when inoculated into seedlings and logs after ten weeks (Figure 4D) and dominated the AOD controls (60 ±4.2% 16S rRNA relative abundance in seedlings and 69 ± 4.1% in logs), while detected at lower relative abundances in SynCom1before and SynCom3before treatments (0.8 ± 1.2% and 1.4 ± 2.5%, respectively, in seedlings and 0.001 ± 0.017 % and 5.8 ± 13.0%, respectively, in logs).

Tree microbiome studies can be restricted due to lack of methods to develop and maintain axenic seedlings (Abdelfattah et al., 2021; Cambon et al., 2025). In our seedlings experiment, 16S rRNA gene profiling of stem microbial cultures did not allow full distinction of SynCom- and endogenous microbiome-derived OTUs across treatments and controls, however, it suggested the presence of some viable SynCom members at the end of the experiment (Figure 4D, Supplementary Table 6).

### AOD-associated bacteria and SynComs induce changes in the oak microbiota

The composition of microbial communities in seedlings inoculated with SynComs was significantly different compared to those in non-inoculated seedlings (Permanova, p-value < 0.05, Supplementary Table 7), suggesting that inoculation of SynComs induced changes in the oak microbiome. ASV differential abundance analysis in seedlings showed that OTUs classified as *Agrobacterium tumefaciens* and *Curtobacterium oceanosedimentum* were enriched in SynCom1before and SynCom3before treatments compared to AOD control (log2 fold change = 26.2 and 27.7, p-value < 0.05). Some contrasting patterns were also observed between different taxa. An OTU in the genus *Streptomyces* was enriched in the AOD control vs Tree Control (log2 fold change = 7.8, p-value < 0.05) but depleted in SynCom1before (log2 fold change = −22.0 and, p-value < 0.05) and an OTU assigned to *Buttiauxella gaviniae* was also enriched in AOD control (log2 fold change = 6.5, p-value < 0.05) and depleted in SynCom3before (log2 fold change = −5.3, p-value < 0.05). Inoculation of SynComs alone seemed to also produce some changes in relative abundance of taxa compared to non-inoculated controls (Extended Data 6 – Abundance Analysis).

Finally, we investigated whether bacterial inoculation negatively affected plant tissue by measuring visibly affected area using the software ImageJ (Extended Data 7, Supplementary Table 5). In the seedlings trial, the size of affected area had a positive correlation with Bg and Gq relative abundances, as well as with the number of gene copies/mL. However, in the logs trial, negative correlations were observed between affected area and pathogen load. Seven out of ten inoculation points in the seedling AOD control evidenced affected areas, while only three out of ten inoculation points were affected in the wound control (inoculated with sterile PBS), which suggests a negative effect caused by AOD-associated bacteria on plant tissue. Nonetheless, SynCom controls (inoculated with SynComs only) also exhibited affected plant tissue, although for SynCom1 and SynCom2 affected areas were, in average, smaller than in treatments inoculated with both SynComs and AOD-associated bacteria. Regarding SynCom treatments, only SynCom1before and SynCom2before AOD exhibited smaller affected areas compared to AOD controls.

The log trial showed differing results. Wound controls exhibited affected areas similar to those in the AOD control (Extended Data 7). This was also the case for SynCom controls (inoculated with SynComs only) and for treatment groups inoculated with SynComs (before or after) plus Bg and Gq. The only exception was SynCom3 control, where lesion areas were smaller than in the AOD control. Overall, this approach suggests that seedlings and logs had different host responses to SynCom inoculation and that some members of the SynCom could potentially cause damage to plant tissue.

## Conclusions

Comprehensive and representative culture collections of microbiota from ecologically important microbiomes are essential to enable phenotypic characterisation of strains, decipher the mechanistic basis of their interactions, and facilitate the rational design of SynComs to understand microbiome dynamics and modulate the microbiome. Here, we generated and screened an extensive microbiota isolate collection from oak trees across Britain, identifying over 300 isolates that suppressed bacterial species associated with stem canker symptoms in oak trees affected by AOD. By assembling and testing SynComs of varying richness *in vitro*, we demonstrated the propensity for SynComs to suppress these bacteria and further validated the protective effect of disease-suppressive SynComs when inoculated *in planta*. The tree stem microbiome is emerging as a critical but underexplored global ecosystem with impacts on tree physiology, carbon sequestration and biogeochemical cycles and is also impacted to diseases caused by canker-causing microbiota. This work highlights the potential to design and apply functionally validated SynComs to engineer the tree stem microbiome for disease-suppression, highlighting opportunities to engineer tree microbiomes for beneficial properties including growth, stress tolerance, disease suppression and chemical cycling.

## Methods

A full overview of the sample collection, preparation and characterization of the microbial culture collection and the selection and identification of bacterial isolates for SynComs can be seen in Extended Data 10.

### Sampling sites and sample collection

We selected twenty sampling sites located within the AOD predisposition zone defined by Brown et al. (2018). Ten of these sites had previous reports of AOD (AOD-sites), and ten sites were located nearby (Near-AOD sites). The remaining ten sites were located outside of the AOD predisposition zone (Far-AOD sites) (Extended Data 10). At each site, leaf, stem and root/rhizosphere samples from five asymptomatic oak trees (*Quercus robur*) were collected for a total of 450 samples (three tissue types x five trees x 30 sites) obtained from 150 oak trees, as described in Downie et al. (2025).

Fully developed leaves were collected from the north and south aspects of the upper portion (25%) of the tree crown. Five leaves from each aspect were selected (total = 10 leaves) and three laminar leaf punches were taken from the bottom, middle and upper part of each leaf using a single-hole punch (30 disks per sample). The remaining leaf tissue was collected in sterile plastic bags and stored at 4 °C. For stem (phloem and sapwood) sample collection, the outer bark of the tree was disinfected with 70% ethanol. A 1.5-cm-arch punch (C. S. Osborne, New Jersey, USA) was placed in between bark cracks at a height of 1.3 m and hammered until reaching the heartwood (approximately 4 cm depth). For rhizosphere sampling, soil cores were obtained using a Dutch soil auger at 1 – 2 m from the base of the tree trunk. Soil cores were placed on disinfected trays and inspected for fine oak roots. Both the fine roots and the portion of soil firmly attached to them were collected. All samples were aseptically separated into two portions, one for microbial isolation (stored at 4 °C until further processing withing 24 h of collection) and one for microbiome analysis (stored at −80 °C in dry ice for further processing).

### Preparation of an oak microbial culture collection

Microbial isolations were conducted at the sampling sites using a dead-air cabinet, unless samples could be transported to the laboratory within 24 h of collection following the methods described in Ordonez et al. (2025). We conducted microbial isolations via (i) agar plating for leaf, stem and rhizosphere and (ii) dilution-to-extinction from leaf and rhizosphere. For agar plating, leaf and rhizosphere samples were homogenized with 50 mL of sterile 1X PBS (PBS, Sigma Aldrich, UK) in a 380-mL portable blender (LaHuko) for 2 min. Cell suspensions were serially diluted up to 10^-1^ and 10^-3^, respectively. Stem samples were surface sterilized in three cycles of immersion in 2.5% sodium hypochlorite for 1 minute followed by 1 min in 1 X PBS, and then air-dried for 30 min. Stem tissue was mixed with 5 mL of 1 X PBS and aseptically ground in mortars and pestles disinfected by 20-min immersion in 10% bleach. Cell suspensions of 100 µL were inoculated on plates containing Reasoner’s 2 agar - R2A (Oxoid, Thermo Fisher Scientific, UK), 10-fold diluted malt extract agar – (Oxoid) and 10-fold diluted potato glucose agar – PGA (Oxoid), all solidified with 2% bacteriological agar, and incubated at ambient temperature for one week.

Bacterial and fungal colonies exhibiting different morphologies were picked using sterile wooden toothpicks and inoculated into 96-well plates containing 200 µL of R2A per well and incubated at room temperature for one week. Isolates were subsequently grown in 200 µL of Tryptone Soy Broth – TSA (Oxoid) in 96-well plates using sterile 96-well pin replicators and incubating for 48 hours. Glycerol stocks were prepared by mixing 100 µL of the liquid cultures with 100 µL of 50% glycerol (Thermo Fisher Chemicals, UK) and stored at −80 °C.

For dilution-to-extinction isolation, we prepared cell suspensions from leaf and rhizosphere samples, and diluted them up to 10^-3^ and 10^-5^, respectively, in a final volume of 60 mL using 10-fold diluted TSB. Cell suspensions of 200 µL were dispensed into each well of three replicates of 96-well plates and incubated for one week at room temperature. A total of 900 96-well plates were produced with the dilution to-extinction-method (three replicates x two tissue types (leaf and rhizosphere) x 150 trees), this is 86,400 wells inoculated in total. Cell suspensions were diluted so that, theoretically, 30 – 50 % of wells contained either a pure culture, or a low-richness community with at least two members. Glycerol stocks were prepared by adding 100 µL of 50% glycerol to 100 µL of culture and stored at −80 °C.

### Single-gene profiling of the oak microbial culture collection

Pools of isolates were prepared by combining 20 µL of glycerol stock from each well of a 96-well plate using Vaccu-Pette/96™ multiwell pipetters (Sigma-Aldrich, UK) and collecting them in 2-mL Eppendorf tubes (volume per plate: ∼2 mL, Supplementary Methods, Section 1). Pools of isolates were centrifuged for 5 min at 10,000 x g and cell pellets were resuspended in 200 µL of nuclease-free water. Genomic DNA was extracted using the boil-prep method, consisting of 5 minutes of incubation at 95 °C, vortexing at 2.5 min incubation.

Two sequencing libraries were prepared for isolates obtained by agar plating and for dilution-to-extinction (Supplementary Methods, Section 2). In both cases, the V4 region of the 16S rRNA gene was amplified using the primers 515F (5’ – GTGBCAGCMGCCGCGGTAA – 3’) and 806R (5’ – GGACTACHVGGGTWTCTAA – 3’) (Caporaso et al., 2011) for identification of bacteria and archaea; and the ITS region was amplified with primers ITS1 (5’ – CTTGGTCATTTAGAGGAAGTAA – 3’) and ITS2 (5’ – GCTGCGTTCTTCATCGATGC – 3’) (Bokulich & Mills, 2013) for identification of fungi. PCR amplification was conducted as described by (Caporaso et al., 2011) (Supplementary Methods, Section 2). For the first library (isolates obtained by agar plating), primer sets contained Illumina adapters, a spacer and an 8-bp barcode and loaded in a MiSeq™ v2 Illumina machine for cluster generation and sequencing by-synthesis in 500 cycles at the Centre for Environmental Biotechnology – CEB (Bangor University, UK) to a depth of approximately 50,000 reads per sample (8.5 Gb raw data). For the second sequencing library (containing isolates obtained by dilution-to-extinction and some repetition samples from agar plating), the primer set only contained 8-bp barcodes and no adaptors and it was sequenced in an Illumina NovaSeq 6000 platform at Novogene (Cambridge, UK), to a depth of approximately 40,000 reads per sample (8 Gb raw data).

### Bioinformatic and statistical analysis of the oak microbial culture collection

Raw sequences from both sequencing runs were demultiplexed using Cutadapt version 3.4 (Marcel, 2011) and processed using the nf-core workflow ampliseq (version 2.7.0) and the reproducible software environments from the Bioconda (Dale et al., 2018) and Biocontainers (Leprevost et al., 2017) projects. This pipeline was executed in Nextflow v23.10.0 (Tommaso et al., 2017). Data quality was evaluated with FastQC (Andrews, 2010) and summarized with MultiQC (Ewels et al., 2016). Primers were trimmed from the raw sequences using Cutadapt. Sequences were processed as one pool with DADA2 (Callahan et al., 2016) to eliminate PhiX contamination, trim reads at 200 bp, filter sequences shorter than 200 bp and sequences with > 2 expected errors, correct errors, merge read pairs, and remove polymerase chain reaction (PCR) chimeras. Taxonomic classification of bacteria and fungi was performed by DADA2 in the ampliseq pipeline using the RDP Naive Bayesian Classifier algorithm (Wang et al., 2007) and the databases ‘Silva 138.1 prokaryotic SSU’ (Quast et al., 2013) for bacterial classification and UNITE v9.0 (Nilsson et al., 2019) for fungal classification.

Output files from the nf-core/ampliseq pipeline (ASV tables, taxonomy tables and associated metadata) were combined into phyloseq objects v1.44.0 (McMurdie & Holmes, 2013) for downstream curation and analysis in R v4.3.1 (R Core Team, 2023). To curate sequencing data sets, we removed ASVs not assigned to phylum level and ASVs classified as chloroplast and mitochondria. If an ASV was present in PCR or DNA negative controls at higher read counts than in true samples, these were removed from the sample. If the ASV in negative controls had lower read counts than in true samples, the number of reads in the negative control was subtracted from the ASV in the sample. Finally, ASVs representing less than 0.1% of the total reads in a sample were also removed (Supplementary Methods, Section 3). Supplementary Table 8 provides a summary of the number of reads per sample types after curation.

Unrestricted Permanova and pairwise restricted Permanova tests using Jaccard distance matrixes (presence/absence-based metric) from the R packages Metacoder v0.3.7 (Foster et al., 2017) and Vegan v2.6-4 (Oksanen et al., 2022), respectively, were used to evaluate differences in the composition of bacterial and fungal cultures obtained from different sites, predisposition zones and isolation methods (Supplementary Methods, Section 3). The proportion of the oak microbiota captured in the microbial culture collection was estimated at different taxonomic levels by comparing 16S rRNA gene and ITS amplicon sequencing datasets against paired datasets from culture-independent analysis of the same oak tissue samples used for microbial isolation (Downie et al., 2025). Heat taxa trees and Venn diagrams were plotted using the R packages Metacoder v0.3.7 and ggVenn package v0.1.10 (Yan, 2023), see Supplementary Methods, Section 3.

### Selection of bacterial isolates and D2E-derived cultures associated with healthy oak trees

Microbial isolates originated from the healthiest oak trees were selected for further characterization, as these may have potential roles in sustaining tree health. The Phenotypic Decline Index (PDI) was used as an indicator of tree health status (Finch et al., 2021). We selected the five 96-well plates with the highest taxonomic richness (based on 16S rRNA sequencing dataset) from the culture collection, obtained from a subset of 13 oak trees exhibiting the lowest PDI in the sampled cohort (n = 150 trees). Plates containing only rhizosphere isolates were excluded, as these could contain microorganisms with limited capacities to thrive in above-ground tree compartments. Isolates and cultures selected were further purified and taxonomically identified.

### *In vitro* screening of the oak microbial culture collection for suppressive activity against AOD-associated bacteria

*B. goodwinii* (FRB 141^T^), the primary causal agent of degraded live tissue in AOD lesion was streaked on nutrient agar and incubated at room temperature for two days. One pure colony was grown overnight in 5 mL nutrient broth at 150 rpm and 28 °C (OD_600_ = 0.4 - 0.6, ∼2.88 x 10^10 CFU/mL). An initial screening was conducted on bacterial lawns prepared (i) by evenly spreading 500 µL of Bg culture on R2A plates and (ii) by inoculating 500 µL of Bg culture in 15 mL of semisolid TSA containing 0.75% (w/v) agar and poured on a plate containing standard TSA. Glycerol stocks of microbial isolates produced by agar plating and contained in 150 96-well plates (n isolates = 14,122) were inoculated on the first type of lawn, while D2E-derived cultures (300 96-well plates) were inoculated on the second type of lawn. We selected 1,089 bacterial isolates/D2E-derived cultures for further confirmation of suppressive activity against *Bg,* using the second lawn approach, which produced clearer inhibition halos.

The suppressive activity of the selected isolates/cultures against the two other bacterial species associated with AOD, *G. quercinecans* (FRB 97^T^) and *R. victoriana* (FRB 225^T^), was also screened using the semisolid agar approach described above. Overnight cultures of these bacteria were first prepared in nutrient broth at 28 °C and 150 rpm. *Gq* and *R. victoriana* cultures were passaged into fresh medium at a 1:10 ratio and incubated for 1.5 and 2 h to reach OD_600_ of 0.1 (9.15 x 10^8) and 0.2 (9.09 x 10^9 CFU/mL), respectively. In all instances screening plates were incubated for five days at room temperature. Inhibition zones were inspected and scored from 1 (very weak inhibition) to 5 (very strong inhibition) based on the size and transparency of the inhibition halo.

### Purification and taxonomic identification of bacterial isolates

Bacterial isolates associated with tree health and pathogen-suppressive bacterial isolates were streaked onto nutrient agar and incubated at room temperature for three days. Single colonies were picked and incubated overnight in 3 mL of TSB at 150 rpm at 28 °C. New glycerol stocks were prepared for further analysis. For suppressive isolates, when more than one colony morphology was observed, each morphotype was separately inoculated into broth and the suppressive activity of each one was validated on semi-solid agar-based assays against *Bg*, *Gq* and *R. victoriana*.

DNA extraction of purified isolates was conducted through boil prep method from 50 µL of bacterial cultures (Supplementary Methods, Section 4). The almost full length of the 16S rRNA gene was amplified using the universal bacterial primers F27 (5′ - AGAGTTTGATCCTGGCTCAG - 3′) and R1492 (5’ - GGWTACCTTGTTACGACTT – 3’) (Heuer et al., 1997). PCR reactions and cycling conditions are described in Supplementary Methods, Section 4. PCR products were Sanger sequenced in forward and reverse directions in the Genewiz sequencing facility (Jiangsu, China).

Sequences were processed with using Geneious Prime® v2023.2.1 (Kearse et al., 2012) (Supplementary Methods, Section 4). Low-quality base calling at the 5’ and 3’ ends were identified and annotated using an error probability limit of 0.01. Forward and reverse sequences (n = 506 isolates) were assembled using De Novo Assembly. In cases where only one direction was successfully sequenced, we used either the forward (n = 4 isolates) or reverse (n = 35 isolates) sequence for further analysis. Basic Local Alignment Search Tool (BLAST) was used for taxonomic identification, using the Megablast algorithm. Consensus sequences were aligned in MEGA 11 (Tamura et al., 2021) using MUSCLE 5.1 (Edgar, 2004) and clustered using the algorithm UPGMA. Phylogenetic trees were created based on the Tamura-Nei model (Tamura & Nei, 1993) and the Neighbour-joining tree-building method (Saitou & Nei, 1987) with 500 Bootstrap iterations.

### Preparation of SynComs and fluorescently labelled AOD-associated bacteria for *in vitro* inhibition assays

One representative isolate of each of the 96 bacterial species (48 pathogen-suppressive and 48 health-associated) identified via Sanger sequencing was selected for preparation of randomly assembled SynComs (Supplementary Methods, Section 5). Bacterial isolates were cultured overnight in 4 mL of TSB at 28°C and 150 rpm. The 96 cultures were randomly distributed across a deep 96-well plate, intercalating pathogen-suppressive and health-associated isolates, and 2 mL per culture were dispensed in an individual well. Isolates were pooled with an 8-channel pipette by rows and columns into sterile reagent reservoirs to prepare SynComs. In total, 40 SynComs composed of either eight (n = 11), 16 (n = 7), 24 (n = 7), 48 (n = 7), 72 (n = 7) or 96 (n = 1) strains were prepared. All members of the SynComs were mixed in equal ratios in final SynCom volume of 500 µL (See Extended Methods). Three technical replicates per SynCom were prepared by pooling strains three times.

SynComs were tested *in vitro* against fluorescently labelled strains *Bg* FRB 186 + pBbE1K + Kan (RFP-Bg) and *Gq* BH1/65b + pBBRI-GFP + Kan (GFP-Gq). The fitness of GFP-Gq was not significantly different than that of the wild-type strain, whilst, RFP-Bg had a higher growth rate than the wild-type strain (Supplementary Methods, Section 6). RFP-Bg and GFP-Gq were streaked in TSA supplemented with kanamycin (10 µg/mL medium) and incubated at room temperature for 48 h. Single colonies were grown overnight in TSB + kanamycin at 28 °C and 150 rpm. Four replicates were prepared, pooled and adjusted to starting OD_600_ of 0.1 for RFP-Bg, and 0.03 for GFP-Gq (minimum OD_600_ at which the fluorescence of each bacterium could be detected).

### *In vitro* evaluation of suppressive activity of SynComs against fluorescently labelled AOD-associated bacteria

For *in vitro* inhibition assays, 190 µL of each fluorescently labelled bacterium were dispensed in one well of a black 96-well microculture plate, and inoculated with 10 µL of a different SynCom (Supplementary Methods, Section 7). We included the following controls: (1) Medium control, containing 190 µL of TSB, (2) Mono-culture control, containing 190 µL of either RFP-Bg or GFP-Gq, (3) SynCom control, containing 190 µL of TSB plus 10 µL of each SynCom, (4) Co-culture control, containing 190 µL of RFP-Bg or GFP-Gq and 10 µL of the wild-type strains of Bg (starting OD_600_ = 0.1) and Gq (starting OD_600_ = 0.03), respectively. The latest was to determine if fluorescently labelled strains could be outcompeted by non-suppressive or neutral strains, such as their wild-type counterparts. All plates were sealed with gas-permeable plate seals (Azenta Life Scicences) and incubated at 28 °C and 150 rpm for 48 h. Relative fluorescence units (RFU) were measured at times 0 h, 24 h and 48 h in a Spectramax M2 fluorescence plate reader (Molecular Devices) at an excitation wavelength of 495 nm and emission of 620 nm for RFP-Bg and at an excitation wavelength of 488 nm and emission of 520 nm for GFP-Gq.

Fluorescence measurements were background- and baseline-corrected (Supplementary Methods, Section 7). Growth curves were produced to calculate the area under the curve (AUC) and estimate the Suppression Index, a quantitative measurement between −100 to 100, using the formula *Suppression Index = (SynCom AUC/Monoculture mean AUC) × 100*, where Monoculture Mean AUC corresponds to the AUC of either RFP-Bg or GFP-Gq, and SynCom AUC corresponds to the AUC of RFP-Bg or GFP-Gq in co-culture with a different SynCom. Negative suppression indexes indicated that RFP-Bg or GFP-Gq did not grow through the experiment and cells died or were unable to express the fluorescent protein. More negative indexes indicate less growth. Positive indexes indicated that RFP-Bg or GFP-Gq were able to grow, although to a lesser extent than the mono-culture controls. The higher the suppression index, the less suppression occurred, with a value of 100 indicating no suppression. Network bipartite plots for the SynComs with highest suppression percentages and the strains contained in them were created using the R package ggraph 2.2.1.

### Statistical analysis of *in vitro* inhibition experiments

Spearman correlation analysis was initially performed to investigate the correlation between the taxonomic richness of SynComs (number of strain members) and the AUC of RFP-Bg and GFP-Gq. Baseline-corrected AUC values (subtraction of fluorescence at time 0 h from times 24 and 48 h) were used for the analysis. Secondly, a generalized linear model (GLM) with a gamma distribution and a log link function was fitted to assess the effect of the relationship between the taxonomic richness of SynComs on the AUC of both fluorescently labelled bacteria. The formula was AUC ∼ Number of SynCom members. To investigate the effect individual strains comprised in the SynComs on the AUC of RFP-Bg and GFP-Gq, a GLM including both the number of strains as well as each bacterial strain as an explanatory variable was applied using the formula ‘AUC ∼ number of SynCom members + bacterial species’ (Supplementary Methods, Section 8). GLMs were performed using the glm function from the R package stats v4.3.1.

### Preparation of SynComs and AOD-associated bacteria for *in planta* inoculation trials

We prepared three SynComs using the same 96 bacterial isolates used for *in vitro* testing: (1) SynCom1, comprising 48 isolates with pathogen-suppressive phenotypes, (2) SynCom2, comprising 48 health-associated isolates, and (3) SynCom3, prepared with 96 strains (48 pathogen-suppressive and 48 health-associated isolates). SynComs were prepared by mixing 2 mL of culture of each isolate and resuspending in 1X PBS to an OD_600_ of 0.6 (Supplementary Methods, Section 9).

We used the type strains Bg FRB 141^T^ and Gq FRB 97^T^ for *in planta* trials (Supplementary Methods, Section 9). Both bacteria were grown overnight in 5 mL of TSB at 28 °C and 150 rpm. Five technical replicates were pooled together and resuspended in Ringer’s solution (OD_600_ = 0.6 for each bacterium). Bg and Gq were mixed in a 10:1 ratio, in order to reflect the strong dominance of Bg over Gq observed in natural conditions by metagenomic studies (Denman et al., 2017).

### Experimental design of *in planta* inoculation trials

A full overview of the experimental design for *in planta* inoculation trials can be found in Supplementary Methods, Section 10. Two independent experiments were conducted using oak seedlings and oak log billets. In both cases, control and treatment groups consisted of 10 inoculation points, either five seedlings with two inoculation points each, or two logs with five inoculation points each. Each experiment had the following controls: (i) Tree control, non-inoculated and non-wounded; (ii) Wounded control, wounded but non-inoculated; (iii) AOD control, inoculated with a Bg and Gq and (iv) SynCom (1, 2 or 3) controls, inoculated one SynCom only, respectively. Six treatments were evaluated, three of them corresponded to inoculation of SynComs 10 days before inoculation of AOD-associated bacteria (SynCom1before, SynCom2before and SynCom3before) and three more corresponding to inoculation of SynComs 10 after inoculation of AOD-associated bacteria (SynCom1after, SynCom2after and SynCom3after). Seedlings and logs were incubated in temperature-controlled rooms for 10 weeks. Our main response variables measuring the load of AOD-associated bacteria were: (1) gene copies/mL of Bg and Gq via qPCR assay and (2) their relative abundance or prevalence via 16S rRNA gene community profiling. In addition, we measured host responses by calculating the area of darkened or degraded tissue in the stem using ImageJ (Schneider et al., 2012).

### Inoculation of SynComs and AOD-associated bacteria in oak seedlings and logs

*In planta* inoculation trials were conducted in temperature-controlled rooms. Six months prior the start of the experiment, watering was progressively reduced in seedlings to predispose them for bacterial infection (See Supplementary Methods, Section 11 for details on plant maintenance and growth). Seedlings were located in randomized blocks in the growth room to account for potential positional effects. Logs were stood on watering trays and the top part of them was sealed with wax.

SynComs and/or AOD-associated bacteria were inoculated into the seedling stems and in the inner bark of logs by wounding the tissue with sterile scalpels and attaching PCR microtubes with bottoms previously removed, directly on the wound. An aliquot of 200 µL of SynComs or AOD-associated bacteria were pipetted inside the PCR tube and sealed with parafilm. After 10 days, seedlings and logs received a second inoculation of either SynComs or AOD-associated bacteria. After the second inoculation, seedlings and logs were incubated for 10 weeks. Samples for qPCR and 16S rRNA gene community profiling were collected at the end of the experiment by swabbing each inoculation point. Pathogen load was determined by (i) qPCR assays (gene copies/mL), and (ii) relative abundance or prevalence of AOD-associated bacteria determined by 16S rRNA gene community profiling. We also collected stem pieces from inoculation points and incubated them in 2 mL of nutrient broth with aerobic static incubation at 28 °C for two days to test the viability of the AOD-associated bacteria and SynComs at the end of the in-planta trials. Growth from stem cultures was analysed via 16S rRNA profiling.

### Quantification of AOD-associated bacteria via qPCR assays

Swabs from inoculation points were washed, filtered to remove large particles and resuspended in nuclease-free water (Supplementary Methods, Section 12). Multiplex real-time PCR assays were performed as described in Crampton et al. (2020), by targeting the gyrB gene of Bg using the primers Bg99F 5’ – CTGGCCGAGCCTGGAAAC-3’ and Bg179R 5’ – AGTTCAGGAAGGAGAGTTCGC – 3’ and the probe Bg124P FAM-5’ – CCAGAATCTCATATTCGAACTCCACCATGTT-BHQ1 – 3’; and the rpoB gene of Gq using the primers Gq284F 3’ – GGCTTTGATAGTGGTGGCC – 3’ and Gq418R 5’-CGTTCCGTTATCACCGTGG – 3’ and the probe Gq342P 5’ – Cy5-AACAGTTCCAGCGCCATTTTCTTCG-BHQ3 – 3’. PCRs and fluorescence detections were performed on a LightCycler480 II instrument (Roche), see Supplementary Methods, Section 12 for detailed qPCR reactions and cycling conditions. DNA standards were prepared from genomic DNA of Bg FRB 141^T^ and Gq FRB 97^T^, extracted with the DNeasy Blood and Tissue kit (Qiagen). DNA standards were normalized to a concentration of 10,000 gene copies/µL and serial dilutions were prepared to create a standard curve. Standards were quantified on a QIAquant-96 (Qiagen) PCR cycler. A standard of 2,000 gene copies of each bacterium was included in each of the qPCR runs for inter-plate calibration of Ct values and determination of the number of gene copies/mL.

### 16S rRNA gene community profiling of oak seedlings and logs

DNA was extracted from swab washes and stem cultures, as well as from SynComs (SynCom inoculum) and the AOD-associated bacteria mix (AOD inoculum) inoculated in seedlings and logs (Supplementary Methods, Section 13). The V4 region of the 16S rRNA gene was amplified using the primer sets mentioned above (Caporaso et al., 2011), containing 8-bp barcodes for sample demultiplexing. PCR reactions, cycling conditions and sequencing library preparation are detailed in Supplementary Methods, Section 13. PCR products were quantified using the Quant-iT™ PicoGreen™ dsDNA Assay (Invitrogen) and pooled to a final concentration per pool of 50 ng for amplicon sequencing. The library was purified using Agencourt AMPure XP Beads (Beckman Coulter) at a bead-to-sample ratio of 0.9:1 (v/v), according to manufacturer’s instructions, and eluted in 100 µL of nuclease-free water. The final sequencing library was sequenced in an Illumina NovaSeq 6000 platform at Novogene (Cambridge, UK), to a depth of approximately 40,000 reads per sample (8 Gb raw data).

### Bioinformatic analysis

Raw sequences were demultiplexed and processed as described above using nf-core workflow ampliseq 2.9.0 (Straub et al., 2020) executed in Nextflow v24.04.2 (Tommaso et al., 2017) (Supplementary Methods, Section 14). ASVs not assigned to phylum level or classified as chloroplast and mitochondria were removed from the whole data set. PCR and DNA negative controls were used to curate the sequencing data sets using the frequency method of the R package ‘decontam’ v1.20.0 (Davis et al., 2018). ASVs with a relative abundance below 2.5% in each individual sample were also removed (Supplementary Methods, Section 14).

ASVs were dereplicated using vsearch v2.14.1 (Rognes et al., 2016) and clustered into OTUs using swarm v3.0.0 (Mahé et al., 2021), see details in Supplementary Methods, Section 14. Briefly, representative sequences of each OTU were used for taxonomic assignment using the RDP Naïve Bayesian Classifier algorithm (Wang et al., 2007) and the database Silva 138.2 prokaryotic SSU. We removed low abundant OTUs with < 10 reads in the entire data set, and low abundant OTUs with taxonomic assignations not expected in the AOD and SynCom inocula from those samples (Supplementary Table 9).

OTUs present in the AOD inoculum were designated as “AOD-derived OTUs” and OTUs obtained in the SynComs inoculum were designated as “SynCom-derived OTUs”. To identify specific OTUs representing the bacterial isolates contained in the SynComs, OTUs were matched against a customized database of the almost full-length sequences of the 16S rRNA gene of the isolates in the SynComs. Details on the construction of the customized database and the alignment of OTUs are explained in Supplementary Methods, Section 14. SynCom-derived OTUs were subsequently tracked across samples in at the end of the *in-planta* experiments to assess the persistence of SynComs. Sequencing data from stem cultures was used to assess the viability of the SynComs and AOD-associated bacteria inoculated.

### Statistical analysis of *in planta* inoculation trials

Spearman corelation analysis were applied to test correlations between (i) gene copies/mL (qPCR) vs relative abundance (16S rRNA sequencing) of each AOD-associated bacterium and (ii) the areas of affected plant tissue vs gene copies/mL and vs relative abundance of pathogens.

Wilcox tests with p-values adjusted using the Benjamini-Hochberg method were applied for an initial assessment of differences between gene copies/mL of Bg and Gq across treatments. Significant differences indicated by Wilcox tests were further investigated using LMMs and GLMMs with gamma distribution to account for potential pseudo-replication effects arisen by multiple inoculation points in the same experimental unit (log and seedling). SynCom treatments were included as fixed effects and either log ID or seedling ID and seedling randomized block number were included as random effects. The relative abundance of Bg and Gq (16S rRNA gene sequencing) across treatments was also modelled with a GLMM with gamma distribution. All mixed-effects models were performed using the functions lmer and glmer from the R package lme4 v1.1-36 using the general formula “pathogen load ∼ treatment + (1|unit ID) + (1|randomized block)” (Supplementary Methods, Section 15).

In the seedlings trial, a differential abundance analysis of ASVs was applied using the DESeq2 v1.40.2 R package (Love et al., 2014) to investigate depleted or enriched OTUs across treatments (Supplementary Methods, Section 15).

## Data availability

The raw data from single-gene community profiling (16S rRNA and ITS) of the oak microbial culture collection, the in-planta trials and sequences of the 16S rRNA gene of bacterial isolates used for SynCom assembly will be available on the European Nucleotide Archive under the project accession PRJEB103756.

## Acknowledgements

This work was funded by the UK Research and Innovation’s (UKRI) Strategic Priorities Fund (SPF) program on Bacterial Plant Diseases (FUTURE OAK project; grant BB/T01069X/1) funded by the Biotechnology and Biological Sciences Research Council (BBSRC), Natural Environment Research Council (NERC), Department for Environmental Food and Rural Affairs (DEFRA), and The Scottish Government. We thank the Forestry Commission’s Technical Services Unit (TSU) staff, especially Liz Richardson, Mark Oram and Alistair MacLeod for assistance with sample collection. We also thank the woodland site owners for permissions to sample oak trees at sites across Britain.

## Author contributions

**Alejandra Ordonez:** Conceptualisation, Fieldwork, Methodology, Investigation, Formal analysis, Data Curation, Writing - Original Draft. **Marine C. Cambon:** Fieldwork, Conceptualisation, Methodology, Investigation, Formal analysis, Data Curation, Writing - Review & Editing. **Jim Downie:** Conceptualisation, Fieldwork, Methodology, Investigation, Writing – Review & Editing, Visualization, Fieldwork, project management, supervision. **Anparasy Kajamuhan:** Methodology, Investigation, Data Curation. **Usman Hussain:** Fieldwork, Investigation, Writing - Review & Editing. **Bridget Crampton:** Methodology, Investigation, Data Curation. **Megan Richardson:** Methodology, Investigation, Data Curation. **Marcus Jones:** Investigation. **Robert Green:** Investigation. **Bethany Pettifor:** Investigation. **Ed Pyne:** Investigation. **Jasen Finch:** Conceptualisation, Resources, Data Curation, Formal analysis. **Carrie Brady:** Investigation, Resources, Writing – Review & Editing. **Sandra Denman:** Conceptualisation, Resources, Methodology, Writing - Review & Editing, Project administration, Funding acquisition. **James E. McDonald:** Conceptualisation, Methodology, Investigation, Writing – Original Draft, Visualization, Supervision, Project administration, Funding acquisition.

