## Extended data for "Large-scale culturing of the tree microbiome enables targeted disease suppression"

**Extended Data 1. Taxonomic composition at the family, order, and class levels of bacterial (A–C) and fungal (D–F) communities detected using culture-dependent and culture-independent approaches.** Single-gene profiling of bacterial and fungal isolates in the culture collection was compared with direct sequencing of the oak microbiome. This comparison revealed an overlap between taxa detected by both types of analysis (Panels ‘Culture-dependent and culture-independent analysis’), as well as taxa identified exclusively by one method (Panels ‘Culture-independent only’ and ‘Culture-dependent only’).

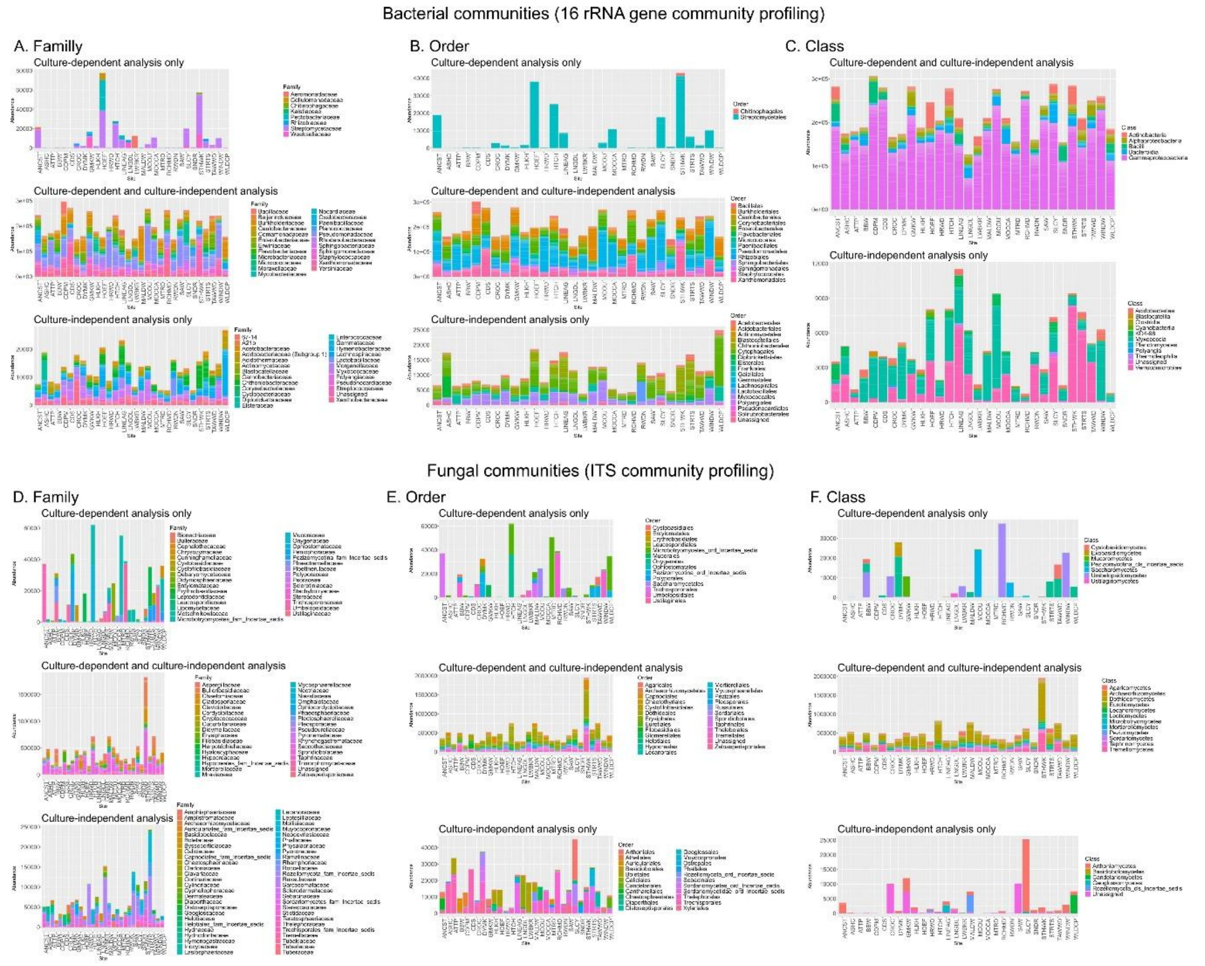

**Extended Data 2. Oak microbial isolates with potential roles supporting tree health.** A) List of species and number of isolates per species with suppressive activity against AOD-associated bacteria (*Brenneria goodwinii, Gibbsiella quercinecans* and *Rahnella victoriana*) and/or associated with healthy oak trees with low phenotypic decline index (PDI). B) Number of species represented by suppressive isolates, tree health-associated isolates and both types of isolates. The suppressive activity of isolates was assessed *in vitro* against each of the AOD-associated bacteria in dual culture-based plate assays. Microbial isolates displaying an inhibition halo against at least one of the AOD-associated bacteria were classified as suppressive. Health-associated isolates were obtained from asymptomatic oak trees with the lowest PDI among the trees sampled (n = 150). Taxonomic assignation was based on Sanger sequences the almost full-length of the 16S rRNA gene.

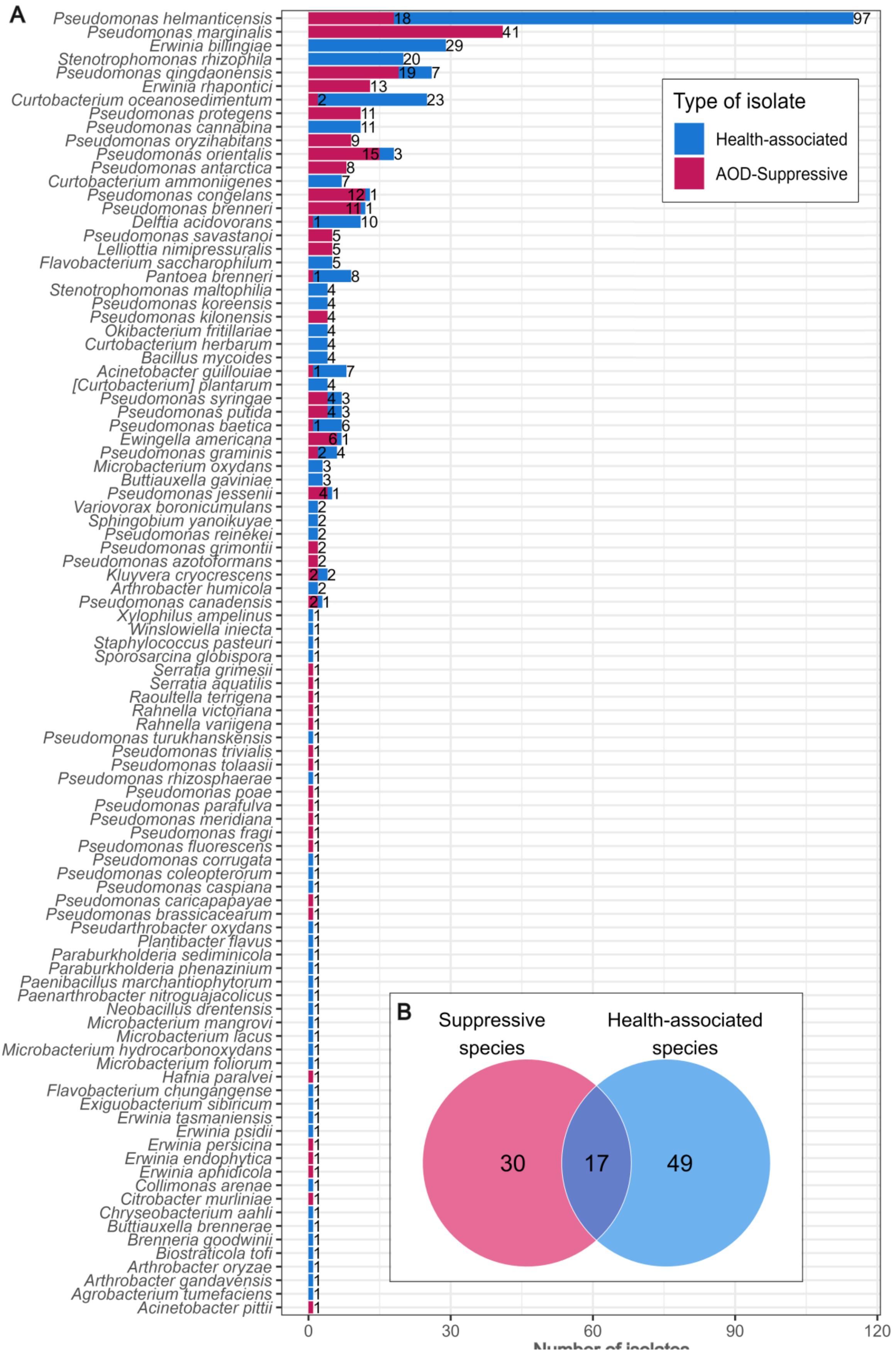
A
Pseudomonas helmanticensis
Pseudomonas marginalis
Erwinia billingiae
Stenotrophomonas rhizophila
Pseudomonas qingdaonensis
Erwinia rhapontici
Curtobacterium oceanosedimentum
Pseudomonas protegens
Pseudomonas cannabina
Pseudomonas oryzihabitans
Pseudomonas orientalis
Pseudomonas antarctica
Curtobacterium ammoniigenes
Pseudomonas congelans
Pseudomonas brenneri
Delftia acidovorans
Pseudomonas savastanoi
Lelliottia nimipressuralis
Flavobacterium saccharophilum
Pantoea brenneri
Stenotrophomonas maltophilia
Pseudomonas koreensis
Pseudomonas kilonensis
Okibacterium fritillariae
Curtobacterium herbarum
Bacillus mycoides
Acinetobacter guillouiae
[Curtobacterium] plantarum
Pseudomonas syringae
Pseudomonas putida
Pseudomonas baetica
Ewingella americana
Pseudomonas graminis
Microbacterium oxydans
Buttiauxella gaviniae
Pseudomonas jessenii
Variovorax boronicumulans
Sphingobium yanoikuyae
Pseudomonas reinekei
Pseudomonas grimontii
Pseudomonas azotoformans
Kluyvera cryocrescens
Arthrobacter humicola
Pseudomonas canadensis
Xylophilus ampelinus
Winslowiella iniecta
Staphylococcus pasteuri
Sporosarcina globispora
Serratia grimesii
Serratia aquatilis
Raoultella terrigena
Rahnella victoriana
Rahnella variigena
Pseudomonas turukhanskensis
Pseudomonas trivialis
Pseudomonas tolaasii
Pseudomonas rhizosphaerae
Pseudomonas poae
Pseudomonas parafulva
Pseudomonas meridiana
Pseudomonas fragi
Pseudomonas fluorescens
Pseudomonas corrugata
Pseudomonas coleopterorum
Pseudomonas caspiana
Pseudomonas caricapapayae
Pseudomonas brassicacearum
Pseudarthrobacter oxydans
Plantibacter flavus
Paraburkholderia sediminicola
Paraburkholderia phenazinium
Paenibacillus marchantiophytorum
Paenarthrobacter nitroguajacolicus
Neobacillus drentensis
Microbacterium mangrovi
Microbacterium lacus
Microbacterium hydrocarbonoxydans
Microbacterium foliorum
Hafnia paralvei
Flavobacterium chungangense
Exiguobacterium sibiricum
Erwinia tasmaniensis
Erwinia psidii
Erwinia persicina
Erwinia endophytica
Erwinia aphidicola
Collimonas arenae
Citrobacter murliniae
Chryseobacterium aahli
Buttiauxella brennerae
Brenneria goodwinii
Biostraticola tofi
Arthrobacter oryzae
Arthrobacter gandavensis
Agrobacterium tumefaciens
Acinetobacter pittii
0
30
60
90
120
Type of isolate
Health-associated
AOD-Suppressive
B
Suppressive species
Health-associated species
30
17
49

**Extended Data 3. Bacterial strains contained in randomly assembled SynComs.** Strains isolated from healthy oak individuals and taxonomically identified using the almost full-length of the 16S rRNA gene and BLAST search were selected to prepare 40 SynComs. The ability of the SynComs to suppress fluorescently labelled strains of the AOD-associated bacteria, *Brenneria goodwinii* (RFP-Bg) and *Gibbsiella quercinecans* (GFP-Gq), was tested *in vitro* in co-culture assays by tracking the targeted strains' fluorescence over 48 hours of growth. A negative suppression index indicated that RFP-Bg or GFP-Gq did not grow through the experiment and cells died or were unable to keep expressing the fluorescent protein. More negative indexes indicate stronger suppression. Positive indexes indicate that RFP-Bg or GFP-Gq were able to grow, although to a lesser extent than the mono-culture controls. The higher the suppression index, the less suppression occurred, with a value of 100 indicating no suppression. Species names contain a suffix indicating the short ID for each isolate used in this study, where 'S' stands for pathogen-suppressive strain and 'H' stands for tree health-associated strain. Number in the y axis correspond to SynCom IDs.

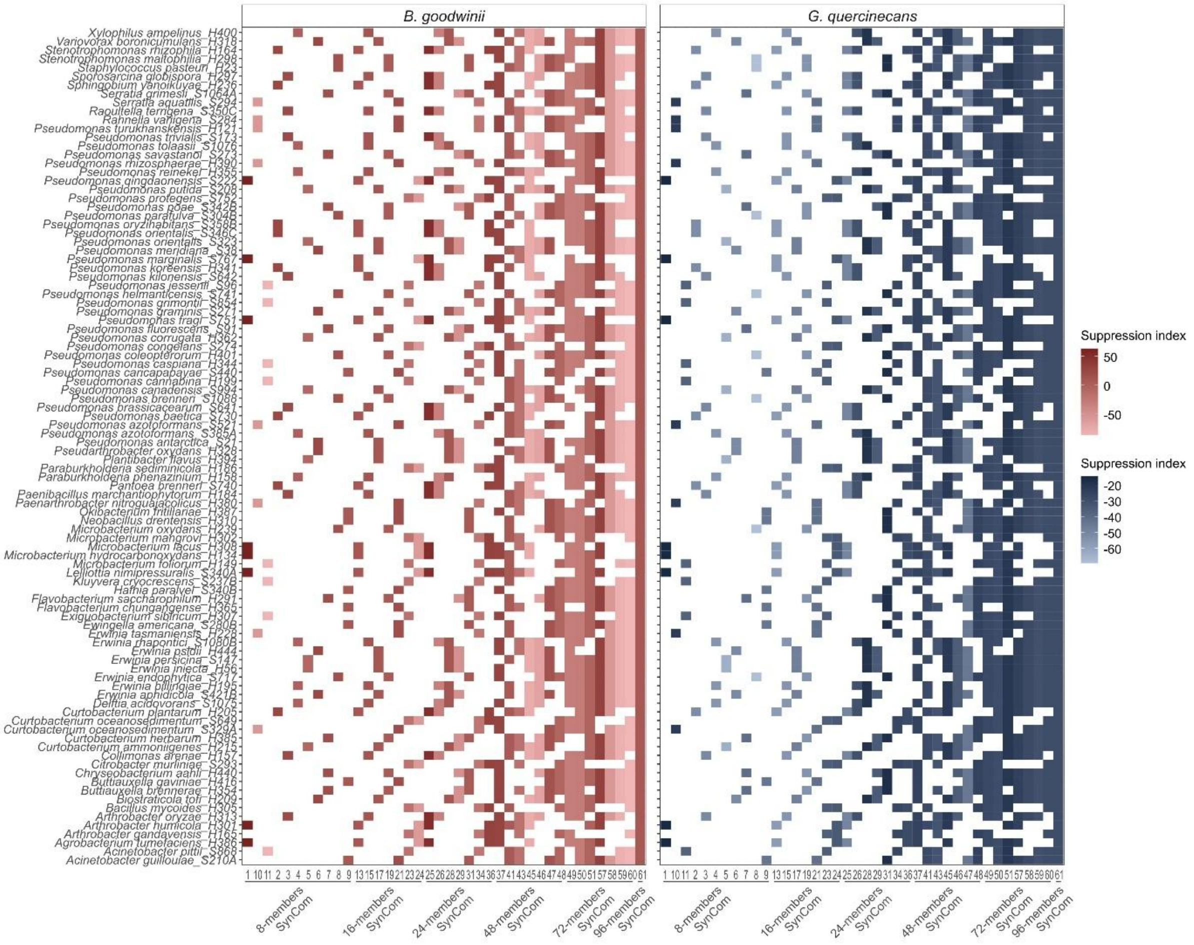

**Extended Data 4. Effect of randomly assembled SynComs on the prevalence (proportion of samples) of AOD-associated bacteria *Brenneria goodwinii* and *Gibbsiella quercinecans* in inoculation trials using oak logs.** Three SynComs, composed of either 48 or 96 strains, were inoculated in logs and seedlings either ten days before or after infection with AOD-associated bacteria. The experimental design included one control inoculated with Bg and Gq (AOD Control) and one controls inoculated with SynComs only (SynComC). The load of AOD-associated bacteria was estimated in swab samples and stem chip cultures via 16S rRNA gene community profiling. Prevalence was determined as the proportion of samples (swabs and stem chip cultures) where Bg and Gq were detected.

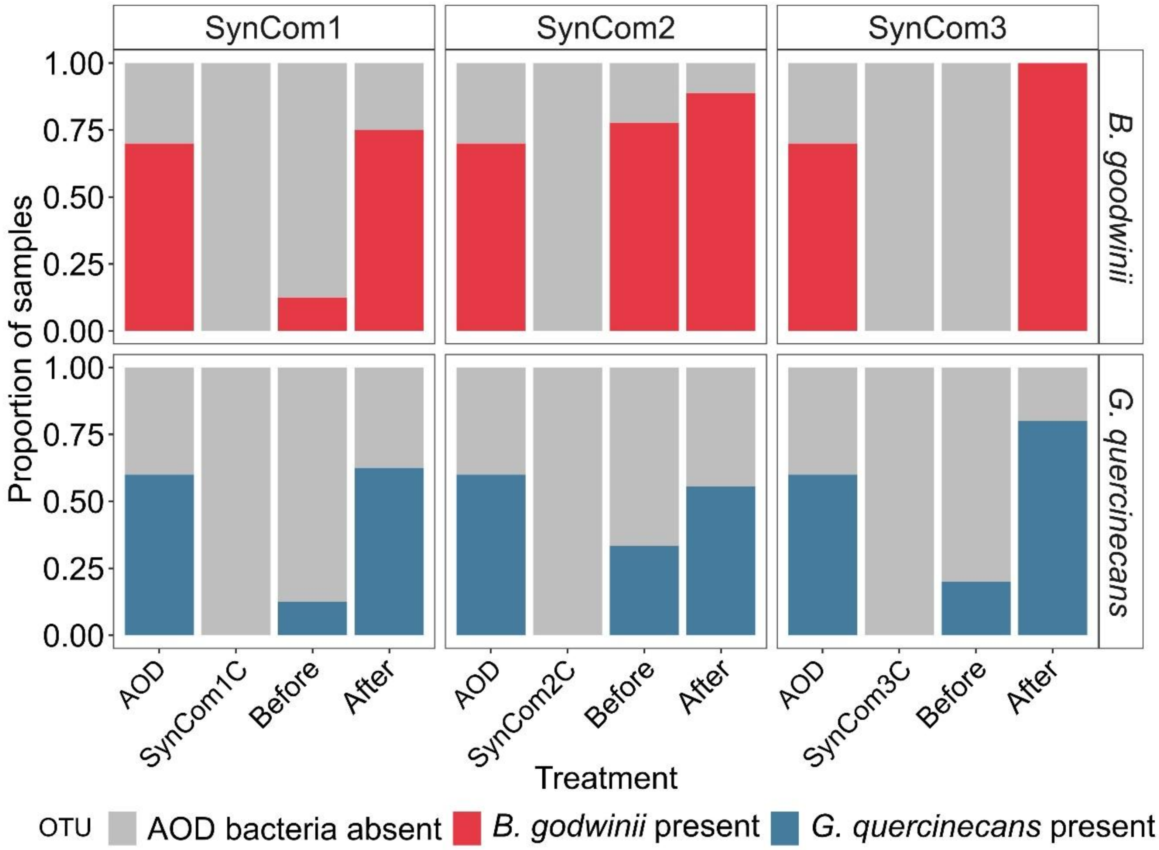

**Extended Data 5. Linear regression between gene copies/mL (qPCR) and number of reads (16S rRNA community profiling) of *B. goodwinii* and *G. quercinecans* in oak seedlings and logs.** Spearman correlation analysis was conducted to assess correlation strength and significance.

A

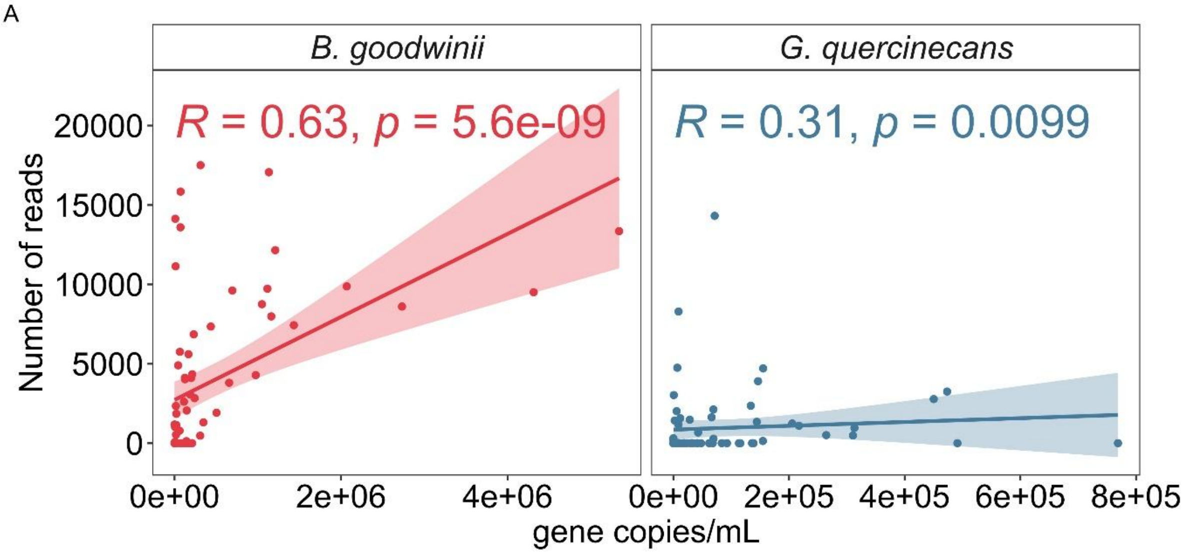

**Extended Data 6. Differential ASV abundance analysis of the oak microbiota in seedlings and logs inoculated with AOD-associated bacteria and randomly assembled SynComs.** Differential ASV abundance analysis was performed for: (A) SynCom1 vs non-wounded, non-inoculated sapling control; (B) SynCom2 vs non-wounded, non-inoculated sapling control; and (C) SynCom3 vs non-wounded, non-inoculated sapling control; (DE) AOD Control vs non-wounded, non-inoculated seedling control; (E) SynCom1before vs AOD Control; and (F) SynCom3before vs AOD Control. Plots show only those OTUs that were significantly enriched or depleted according to the abundance analysis (p-values < 0.05). Sequences identified as *B. goodwinii* and *G. quercinecans* were confirmed using the BLAST algorithm and are marked with a star symbol. OTUs present in the SynCom inoculum and matched to almost full-length 16S rRNA gene sequences are labelled with an '-S' suffix.

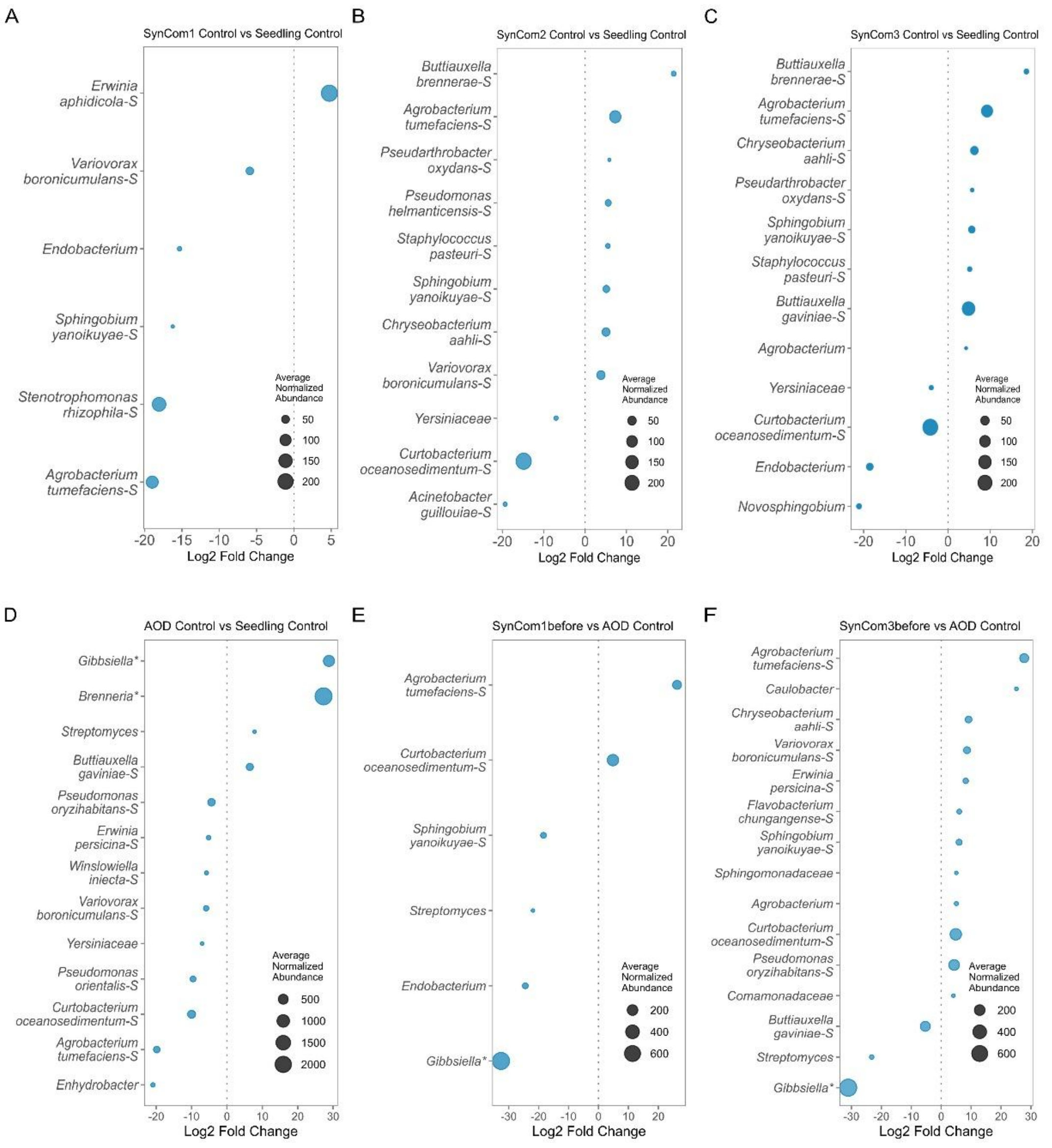

**Extended Data 7. Area of affected plant tissue in oak seedlings and logs inoculated the AOD-associated bacteria (*Brenneria goodwinii* and *Gibbsiella quercinecans*) and/or with randomly assembled SynComs comprising strains obtained from the cultivable oak microbiota.** A - B) Linear regressions between the lesion area of oak seedlings and logs and the number of gene copies/mL (A) and relative abundance (B) of Bg and Gq. A Spearman correlation analysis was conducted to assess the strength and significance of the correlation. C - D) Measurements of lesion area in seedlings and logs, respectively. Wilcox rank-sum tests with Benjamini-Hochberg p-value adjustment were used to compare SynCom treatments against the AOD control**.**

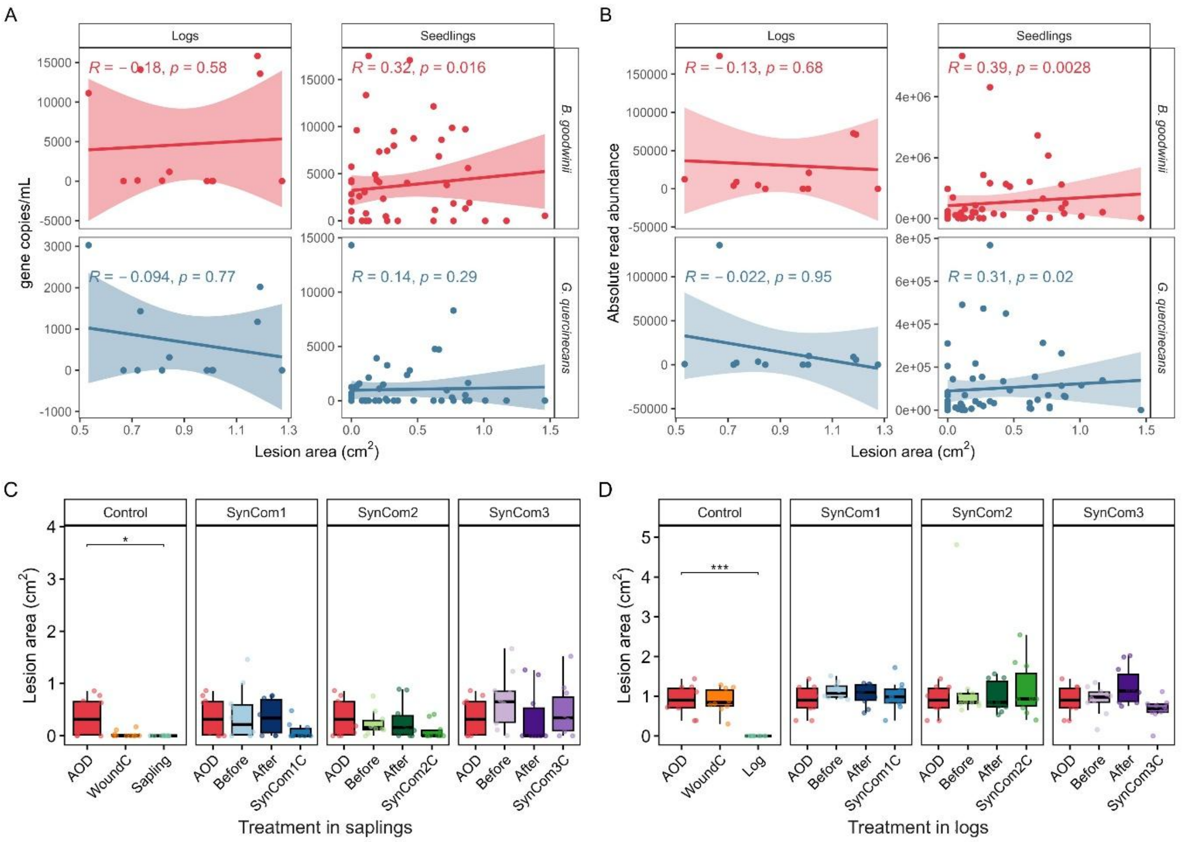

**Extended Data 8. Summary of model results for SynCom effects on the concentration and relative abundance of AOD-associated bacteria in seedlings and logs.** Three SynComs were inoculated either ten days before or after inoculation with AOD-associated bacteria (*Brenneria goodwinii* and *Gibbsiella quercinecans*) in oak seedlings and logs. The concentration of each bacterium was calculated by qPCR assays as gene copies/mL and the relative abundance was estimated by 16S rRNA gene community profiling in swab samples from seedlings and in swab and stem chips cultures from logs. Response variables were obtained from 10 inoculation points (2 inoculation points per seedling and 5 inoculation points per log). The dataset was separated as treatments 'before' and 'after' for each bacterium and for each trial, and a different model was run for each subset. SynCom treatment (SynCom 1, 2 or 3 before/after inoculation with AOD-associated bacteria) were included as fixed effects; and log or seedling unit and seedling block were included as random effects.

| ***Response variable*** | ***Model*** | ***Trial*** | ***Treatment*** | ***Brenneria goodwinii*** | | | | ***Gibbsiella quercinecans*** | | | |
|---|---|---|---|---|---|---|---|---|---|---|---|
| | | | | **Esti-mate** | **Std. error** | **t-value** | **p-value** | **Esti-mate** | **Std. error** | **t-value** | **p-value** |
| *Gene copies/mL* | Linear mixed-effect model (LMM) | Seedlings | SynCom1before | -2.00 | 1.47 | -1.36 | 0.19 | -541 | 1.66 | 0.03 | **3e-03** |
| | | | SynCom2before | 0.82 | 1.47 | 0.56 | 0.59 | -1.97 | 2.00 | -0.98 | 0.34 |
| | | | SynCom3before | 0.03 | 1.47 | 0.02 | 0.99 | -2.77 | 2.00 | -1.38 | 0.19 |
| | | | SynCom1after | 0.42 | 0.86 | 0.49 | 0.63 | 0.04 | 0.95 | 0.05 | 0.96 |
| | | | SynCom2after | -0.69 | 0.86 | -0.80 | 0.43 | -1.62 | 0.95 | -1.70 | 0.12 |
| | | | SynCom3after | -0.11 | 0.86 | -0.13 | 0.90 | -0.37 | 0.95 | -0.39 | 0.70 |
| | | Logs | **SynCom1before** | -5.51 | 1.58 | -3.49 | **9e-04** | -5.24 | 1.73 | -3.02 | **0.02** |
| | | | SynCom2before | -2.87 | 1.58 | -1.81 | 0.07 | -1.21 | 1.68 | -0.80 | 0.48 |
| | | | **SynCom3before** | -3.30 | 1.58 | -2.09 | **0.04** | -4.40 | 1.73 | -2.54 | **0.04** |
| | | | SynCom1after | -0.77 | 1.15 | -0.67 | 0.51 | 1.71 | 1.80 | 0.95 | 0.40 |
| | | | SynCom2after | -2.07 | 1.15 | -1.80 | 0.08 | -0.49 | 1.80 | -0.27 | 0.80 |
| | | | SynCom3after | -1.02 | 1.15 | -0.89 | 0.38 | 0.23 | 1.80 | 0.13 | 0.90 |
| | Generalized linear model (GLM) | Seedlings | SynCom1before | -0.19 | 0.32 | -0.58 | 0.56 | -0.67 | 0.76 | -0.88 | 0.38 |
| | | | SynCom2before | 0.07 | 0.32 | 0.21 | 0.83 | -0.19 | 0.76 | -0.26 | 0.80 |
| | | | SynCom3before | 0.00 | 0.32 | 0.01 | 0.99 | -0.29 | 0.76 | -0.38 | 0.71 |
| | | | SynCom1after | 0.04 | 0.09 | 0.40 | 0.69 | 0.00 | 0.32 | 0.01 | 0.99 |
| | | | SynCom2after | -0.06 | 0.09 | -0.63 | 0.53 | -0.16 | 0.32 | -0.50 | 0.62 |
| | | | SynCom3after | -0.02 | 0.09 | -0.21 | 0.84 | -0.03 | 0.32 | -0.11 | 0.91 |
| | | Logs | SynCom1before | -0.81 | 0.83 | -0.98 | 0.33 | -1.09 | 0.97 | -1.12 | 0.26 |
| | | | SynCom2before | -0.34 | 0.83 | -0.41 | 0.68 | -0.17 | 0.97 | -0.17 | 0.86 |
| | | | SynCom3before | -0.40 | 0.83 | -0.49 | 0.63 | -0.82 | 0.97 | -0.84 | 0.40 |
| | | | SynCom1after | -0.08 | 0.43 | -0.19 | 0.85 | 0.20 | 0.57 | 0.34 | 0.73 |
| | | | SynCom2after | -0.23 | 0.43 | -0.55 | 0.58 | -0.06 | 0.57 | -0.11 | 0.91 |
| | | | SynCom3after | -0.11 | 0.43 | -0.26 | 0.80 | 0.03 | 0.57 | 0.05 | 0.96 |
| *Relative abundance based on 16S rRNA gene community profiling* | Generalized linear mixed-effects model (GLMM) | Seedlings | **SynCom1before** | -2.03 | 0.87 | -2.33 | **0.02** | -8.11 | 0.71 | -11.37 | **1e-16** |
| | | | **SynCom2before** | 0.50 | 0.80 | 0.63 | 0.53 | -2.84 | 0.65 | -4.37 | **1e-05** |
| | | | **SynCom3before** | 0.43 | 0.91 | 0.47 | 0.64 | -8.11 | 0.75 | -10.87 | **1e-16** |
| | | | **SynCom1after** | -0.05 | 0.90 | -0.05 | 0.96 | -4.03 | 1.92 | -2.10 | **0.04** |
| | | | **SynCom2after** | -0.12 | 0.78 | -0.16 | 0.87 | -5.28 | 1.74 | -3.03 | **2e-03** |
| | | | **SynCom3after** | -0.02 | 0.80 | -0.03 | 0.98 | -4.39 | 1.75 | -2.52 | **0.01** |
| | | Logs | **SynCom1before** | -1.91 | 0.82 | -2.32 | **0.02** | -2.78 | 1.02 | -2.74 | **6e-03** |
| | | | **SynCom2before** | -0.37 | 0.80 | -0.46 | 0.65 | -2.06 | 0.98 | -2.09 | **0.03** |
| | | | **SynCom3before** | -8.12 | 0.95 | -8.60 | **2e-16** | -1.14 | 1.17 | -0.97 | 0.33 |
| | | | SynCom1after | . | 0.64 | -0.16 | 0.88 | -0.77 | 0.88 | -0.88 | 0.38 |
| | | | SynCom2after | 0.36 | 0.62 | 0.58 | 0.56 | -0.86 | 0.85 | -1.00 | 0.32 |
| | | | SynCom3after | -0.12 | 0.74 | -0.16 | 0.88 | -0.53 | 1.02 | -0.52 | 0.60 |

**Extended Data 9. Q-Q and Fitted vs Residuals plots from Linear Mixed Models (LMM) of *Brenneria goodwinii* (A - D) and *Gibbsiella quercinecans* (E - H) gene copies/mL in oak seedlings and logs treated with SynComs before or after pathogen inoculation.** Three SynComs were inoculated either ten days before or after inoculation with AOD-associated bacteria (*B. goodwinii* and *G. quercinecans*). The concentration of each bacterium was calculated by qPCR assays as gene copies/mL. The dataset was separated as treatments 'before' and 'after' for each bacterium and for each trial. An LMM was applied for each subset of data. SynCom treatment (SynComs before or SynComs after) were included as fixed effects; and log or seedling unit and seedling block were included as random effects.

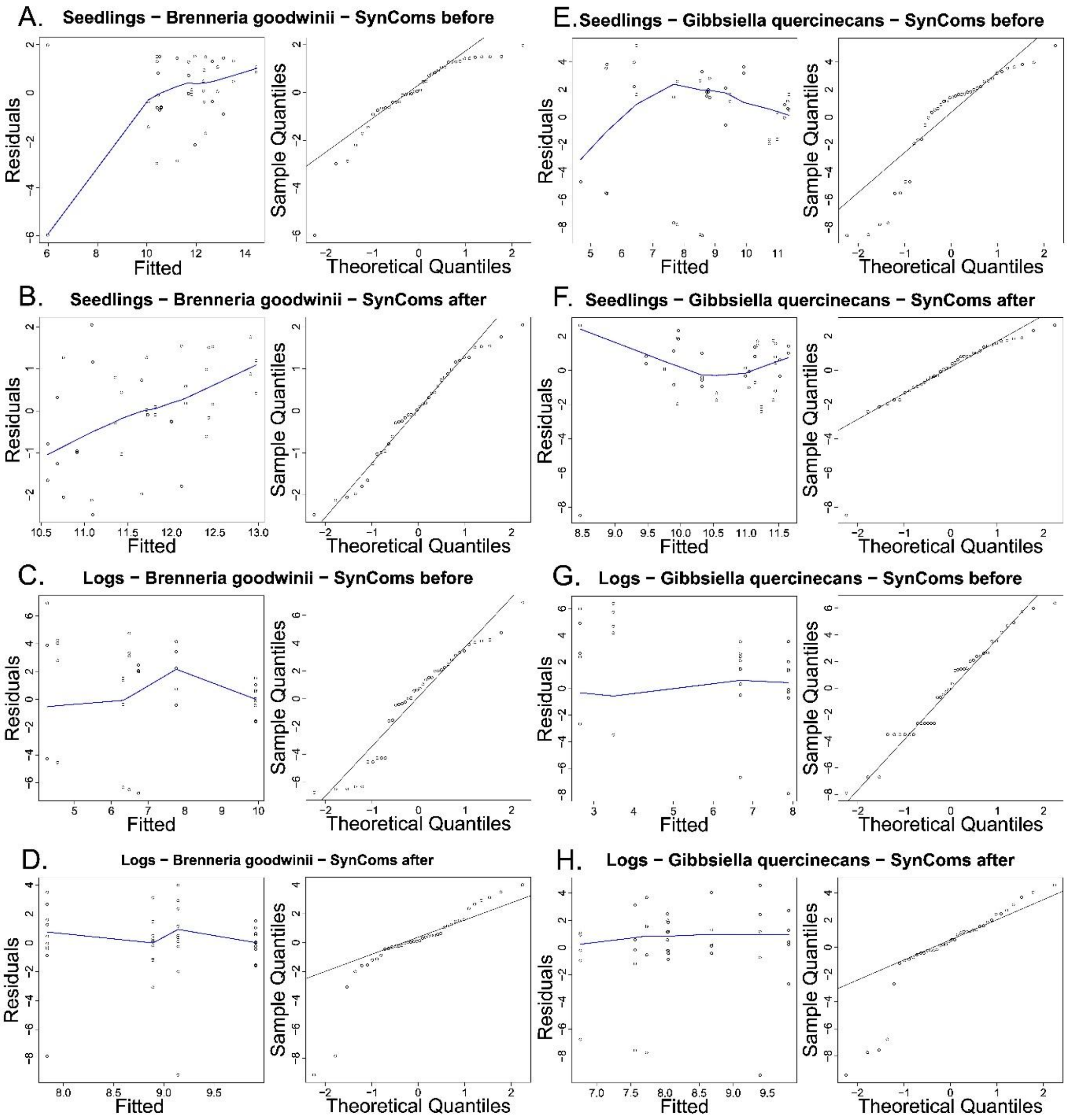

**Extended Data 10. Workflow for establishing a microbial culture collection representative of the oak microbiota and selection of oak microbiota with potential roles in supporting tree health.** A – B) Leaf, stem and root/rhizosphere samples were collected from 150 oak trees located in 30 sites across Great Britan (n = 450). C) A microbial culture collection from oak tissue samples was produced by (i) Agar plating and (ii) dilution-to-extinction. Leaf, stem and root/rhizosphere samples were serially diluted and plated onto three types of agars. Colonies were picked and glycerol stocks were prepared in 96-well plates. One plate containing isolates from the three tissue types was obtained per tree (n = 150). Leaf and root/rhizosphere samples were diluted and dispensed in 96-well plates, 3 replicates each (dilution-to-extinction). Glycerol stocks were prepared from each 96-well plate (n = 900: 150 trees x 2 tissue types x 3 replicates). D) The oak microbial culture collection was characterized by single-gene profiling of the 16S rRNA gene and ITS for bacterial and fungal identification, respectively (culture-dependent sequencing). Isolates contained in each 96-well plates were pooled into one sample. All samples obtained by agar plating (n = 150) and approximately one-third of the plates obtained by dilution-to-extinction (n = 360) were sequenced. E) A parallel microbiome wide association study was conducted on the same oak tissue samples used for microbial isolation (Downie et al., 2025). Tissue samples were sequenced directly after collection using single-gene community profiling of the 16S rRNA gene and ITS for bacterial and fungal identification, respectively (culture-independent sequencing). F) The microbial culture collection was screened in agar-based inhibition assays against the AOD-associated bacteria *B. goodwinii, G. quercinecans* and *R. victoriana*. The phenotypic decline index (PDI) of the oak trees samples was used as an estimation of health status of the tree. Microculture plates with isolates obtained from the trees exhibiting the lowest PDI and with highest taxonomic richness according to sequencing data were selected. Isolates were purified and the full length of the 16S rRNA gene was sequenced by Sanger technology.

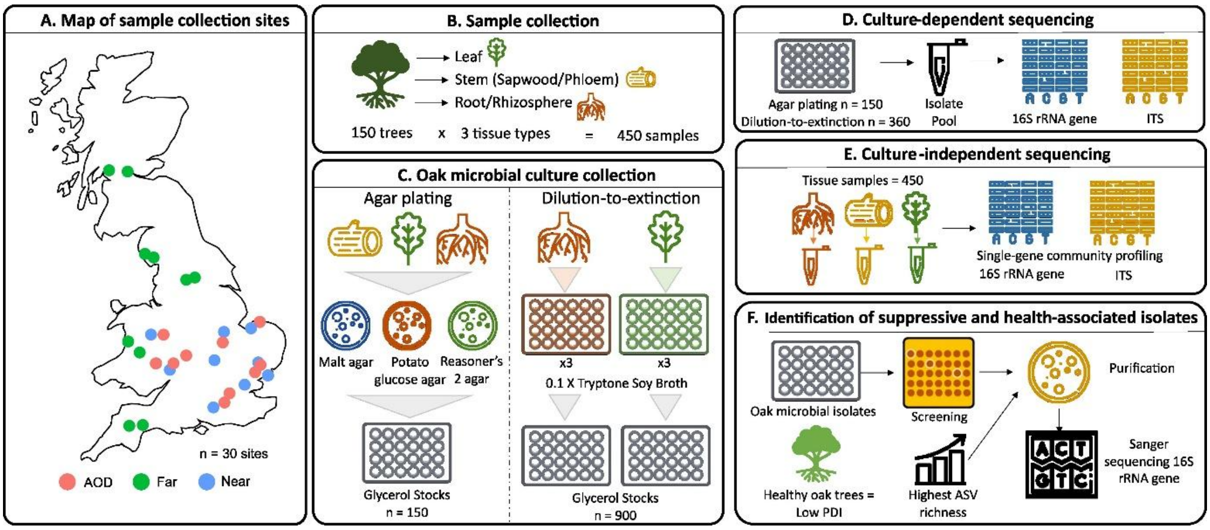
